# Brain-Informed Fine-Tuning for Improved Multilingual Understanding in Language Models

**DOI:** 10.1101/2025.07.07.662360

**Authors:** Anuja Negi, Subba Reddy Oota, Anwar O Nunez-Elizalde, Manish Gupta, Fatma Deniz

## Abstract

Recent studies have demonstrated that fine-tuning language models with brain data can improve their semantic understanding, although these findings have so far been limited to English. Interestingly, similar to the shared multilingual embedding space of pretrained multilingual language models, human studies provide strong evidence for a shared semantic system in bilingual individuals. Here, we investigate whether fine-tuning language models with bilingual brain data changes model representations in a way that improves them across multiple languages. To test this, we fine-tune monolingual and multilingual language models using brain activity recorded while bilingual participants read stories in English and Chinese. We then evaluate how well these representations generalize to the bilingual participants’ first language, their second language, and several other languages that the participants are not fluent in. We assess the fine-tuned language models on brain encoding performance and downstream NLP tasks. Our results show that bilingual brain-informed fine-tuned language models outperform their vanilla (pretrained) counterparts in both brain encoding performance and most downstream NLP tasks across multiple languages. These findings suggest that brain-informed fine-tuning improves multilingual understanding in language models, offering a bridge between cognitive neuroscience and NLP research. We make our code publicly available. ^2^

## 1 Introduction

Recent research has demonstrated that representations extracted from text-based Transformer language models can be used to accurately predict human brain activity evoked during language processing, suggesting parallels between artificial and brain language representations (Wehbe et al., 2014b; Jain & Huth, 2018; Toneva & Wehbe, 2019; Schrimpf et al., 2021; Caucheteux & King, 2022; Goldstein et al., 2022; Karamolegkou et al., 2023; Oota et al., 2025). Although these models accurately predict patterns of brain activity, they have not been originally pretrained to do so. Several previous studies hypothesized that fine-tuning language models with brain data can lead to representations that are better aligned with brain activity (Schwartz et al., 2019; Moussa et al., 2025; Vattikonda et al., 2025). However, these efforts have largely focused on monolingual language models and monolingual brain data, usually trained and evaluated only in English. This overlooks the widespread prevalence of bilingualism and multilingualism in the human population (Grosjean, 2024). The limitation is particularly significant given recent neuroscientific evidence revealing that bilingual individuals have shared semantic representations across languages (Chen et al., 2024b; Francis, 2005), hinting at a shared component for the processing of different languages in the human brain (de Varda et al., 2025). Complementary NLP research has shown that multilingual language models operate in a shared, language-agnostic conceptual space (Wendler et al., 2024; Schut et al., 2025). This raises the question of whether *fine-tuning language models with bilingual brain data can elicit multilingual capabilities in language models*.

In this study, we use brain recordings of bilingual participants reading the same naturalistic stories in English and Chinese, from (Chen et al., 2024b), and investigate whether fine-tuning language models with brain data from bilingual participants can elicit multilingual capabilities in language models. We first introduce a novel, end-to-end brain-informed fine-tuning pipeline (as shown in Fig. 1), which can be trained using data from the whole-brain, or from language- or semantically-selective brain regions. Second, we use brain recordings of bilingual participants to fine-tune two monolingual language models (BERT for English and Chinese (Devlin et al., 2019)) and four multilingual language models (mBERT (Devlin et al., 2019), XLM-R (Conneau et al., 2020), XGLM (Lin et al., 2022), LLaMA-3.2 (Touvron et al., 2023)). Third, we investigate how monolingual and multilingual language models change after fine-tuning with bilingual brain data. We evaluate this along two axes: brain encoding and downstream NLP task performance. To assess generalization across languages, we evaluate the brain-informed fine-tuned language models on downstream NLP benchmarks not only in the models’ fine-tuned language but also in the bilingual participants’ other language (cross-language transfer between known languages) and several other languages that the participant is not fluent in (and also not used in fine-tuning; referred to as ‘unseen languages’), e.g., German and French. Fine-tuning is done separately for each participant, and we assess encoding performance both within the same participant and across the other participants.

**Figure 1:**
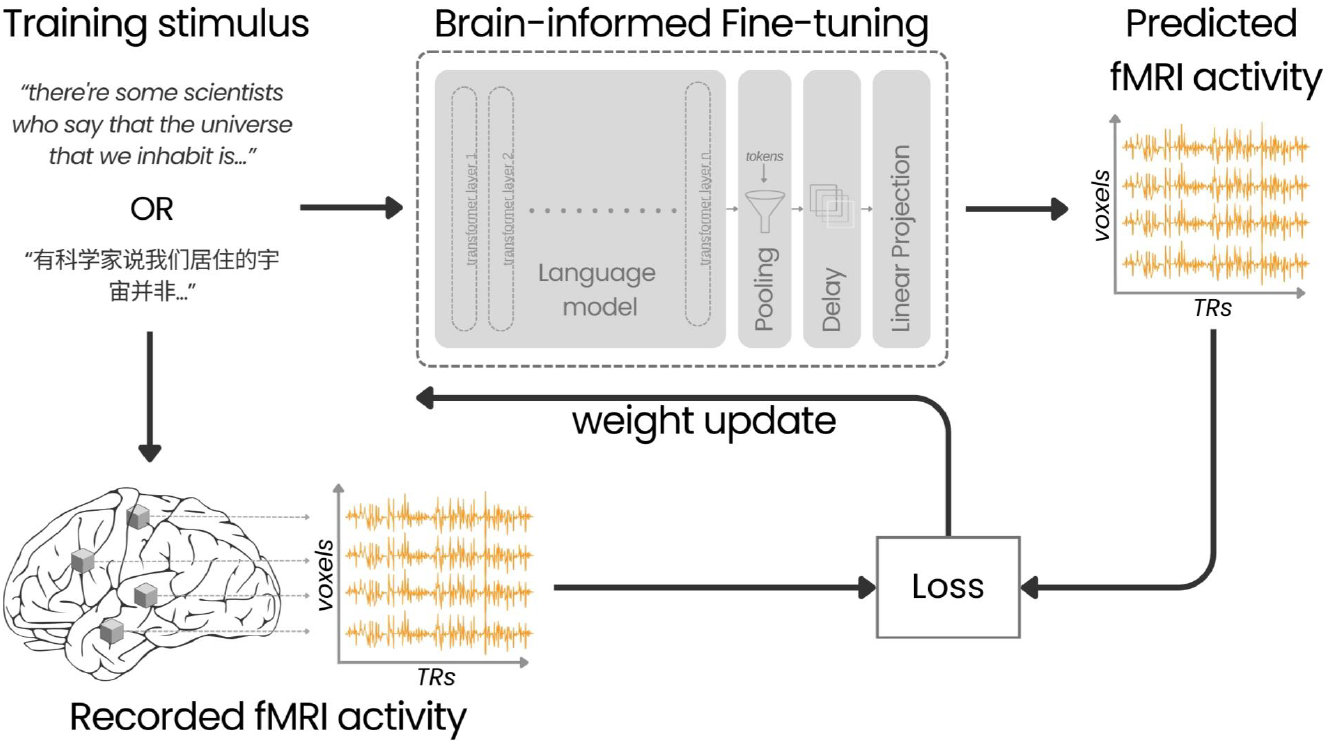
Brain-informed fine-tuning pipeline. Participants read naturalistic stories in English or Chinese while fMRI responses were recorded. The corresponding transcripts were fed into a pretrained language model. Representations from the final token of the last hidden layer were downsampled and temporally delayed to align with the fMRI acquisition. These were then projected to voxel space via a linear layer to predict brain activity. The loss between predicted and recorded responses was backpropagated through all layers for full model fine-tuning.

**Figure 2:**
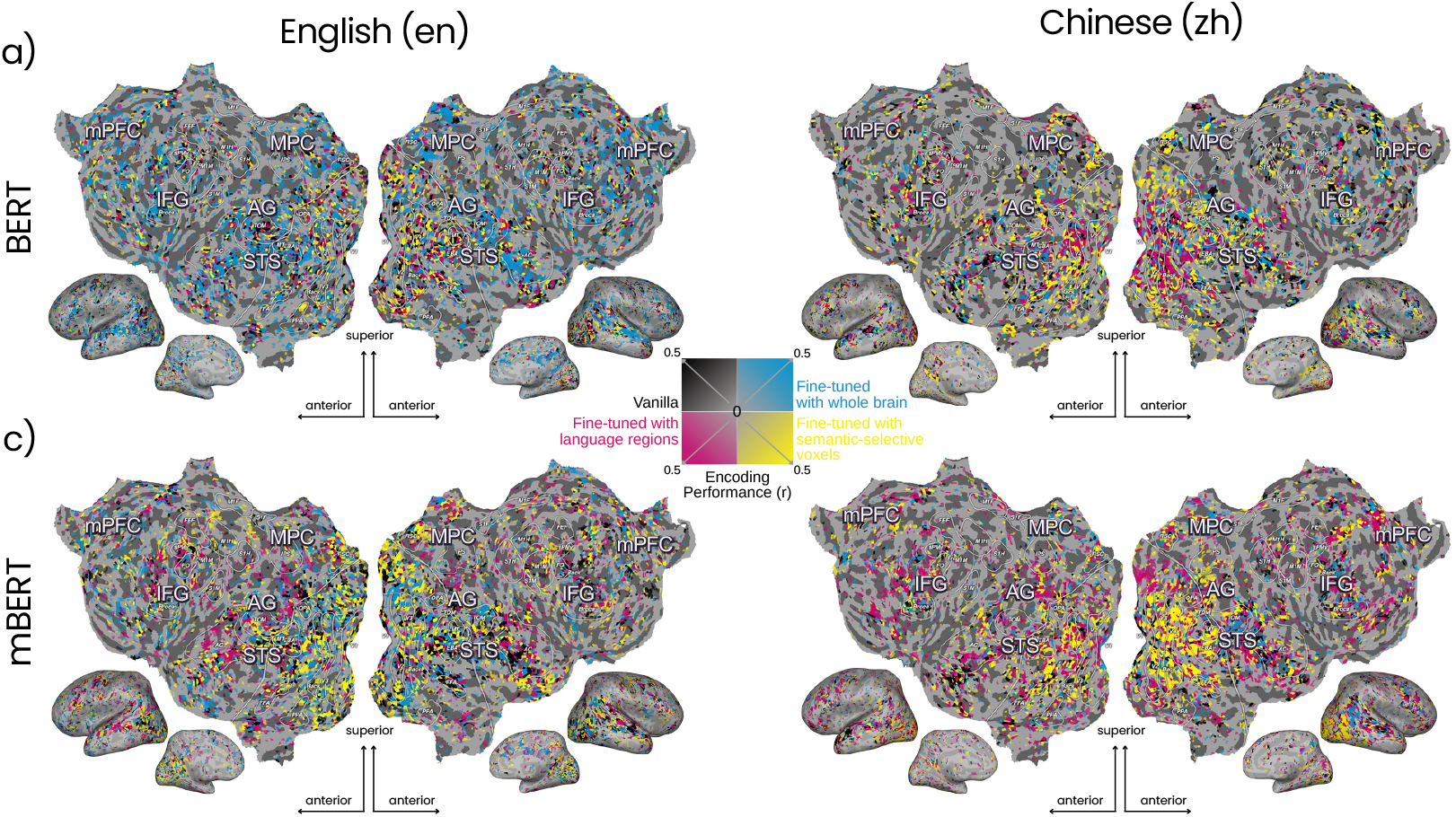
Brain encoding performance before and after brain-informed fine-tuning. We evaluated the effect of bilingual brain-informed fine-tuning using voxelwise encoding models (VEMs; see Methods). VEMs were trained using representations from both the vanilla (pretrained) model and their bilingual brain-informed fine-tuned variants (using whole-brain or using language, or semantically-selective brain regions). Flattened cortical surface for one participant is shown. Each voxel on the cortical surface is colored according to which model variant achieved the best encoding performance (*r*) for (a) BERT and (b) mBERT with English (left) and Chinese (right) brain data, respectively. Each point corresponds to a voxel that was consistently well predicted (*r*>0.1) across models. Voxel colors reflect the best-performing model: black for vanilla, blue for fine-tuned with whole-brain, magenta for fine-tuned with language, and yellow for semantically-selective brain regions. Across both languages and model types, bilingual brain-informed fine-tuning consistently yields better encoding performance than its vanilla counterpart. Results for other participants are in Appendix F.2.

Brain-informed fine-tuning with bilingual brain data reveals several key conclusions: (1) Brain encoding performance improves after fine-tuning, even when evaluated on participants not used in the training set. This suggests that the brain bias introduced is not specific to a particular participant but rather reflects a shared component of bilingual representations. (2) Both monolingual and multilingual language models show improved performance on downstream NLP tasks in the fine-tuned language, the participant’s other language, and in several other unseen languages, indicating that brain-informed fine-tuning elicits generalizable semantic structure not tied to any one language. (3) To ascertain whether our results are specific to fine-tuning with bilingual brain data, and not a general outcome of fine-tuning language models with brain data, we performed the same analyses using brain data from monolingual individuals. Results suggest that the observed effects are indeed driven by bilingual brain representations. These findings suggest that bilingual brain-informed fine-tuning improves multilingual understanding in text-based language models. Our results contribute to the alignment between brain and artificial multilingual language representations, offering insights into the development of brain-inspired multilingual NLP systems.

We make the following contributions: (1) To the best of our knowledge, this is the first study to perform brain-informed fine-tuning using bilingual brain data, applying it to both monolingual and multilingual language models. (2) We introduce a novel brain-informed fine-tuning pipeline that explicitly models the temporal component of brain activity. This contrasts with previous brain-based fine-tuning studies, where these are implemented as preprocessing steps before fine-tuning the model. (3) We evaluate the performance of brain-informed fine-tuned monolingual and multilingual language models on downstream NLP tasks in both English and Chinese. The code is publicly available ^2^.

## 2 Related Work

### Fine-tuning of language models with naturalistic brain data

Our work builds on the brain-tuning approach introduced by Schwartz et al. (2019); Moussa et al. (2025); Vattikonda et al. (2025), which fine-tunes pretrained Transformer-based language models using brain data to integrate brain-relevant information. Schwartz et al. (2019) demonstrated improved brain encoding and NLP task performance using brain data from monolingual English readers, while Moussa et al. (2025); Vattikonda et al. (2025) extended this to speech-based models to enhance semantic representations. Our study complements these by exploring bilingual brain-informed fine-tuning and analyzing how monolingual and multilingual models change when trained with bilingual brain data.

### Multilingual language models and brain alignment

Our work aligns with a growing body of research examining the alignment between human brain activity and language models. Several studies have shown that text-based models can predict brain responses to written and spoken stimuli with high accuracy (Wehbe et al., 2014a; Jain & Huth, 2018; Toneva & Wehbe, 2019; Deniz et al., 2019; Abdou et al., 2021; Toneva et al., 2022; Antonello et al., 2021; Oota et al., 2022; Aw & Toneva, 2023; Oota et al., 2024b; Lamarre et al., 2022; Chen et al., 2024a). Recent efforts have extended this to multilingual Transformer-based models using brain data from tasks in multiple languages (de Varda et al., 2025), though most studies remain monolingual in design, with the exception of Chen et al. (2024a), who explored bilingual processing using similar English and Chinese stimuli. Our study supplements this by investigating bilingual brain alignment through brain-informed fine-tuning and its impact on downstream NLP tasks in both languages. Detailed related work is in Appendix A.

## 3 Methodology

### 3.1 Naturalistic Brain Imaging Dataset

#### Bilingual fMRI dataset

Blood oxygen level-dependent (BOLD) responses were collected using fMRI from six bilingual participants (three males, three females), all native speakers of Mandarin Chinese and fluent in English as a second language. Participants read 11 narrative stories from The Moth Radio Hour (Huth et al., 2016), presented word-by-word in separate scanning sessions for English (en) and Chinese (zh). This dataset is taken from Chen et al. (2024b). Each story is 10-15 minutes long. Of the 11 stories, seven were used for fine-tuning the language models (covering 2756 TRs ^3^), three were used for training voxelwise encoding models (1117 TRs), and one story was reserved for testing (291 TRs). The same set of stories were used in each language to ensure comparability across languages.

#### Monolingual fMRI dataset

To test whether the effects of brain-informed fine-tuning are specific to bilingual brains, we compare models fine-tuned with fMRI data from three English-monolingual participants (all male; one from Deniz et al. (2019) and two from LeBel et al. (2023)). The participants read or listened to the same English narrative stories under similar experimental designs.

This allows direct comparison between models fine-tuned with bilingual and monolingual brain data. If similar improvements are seen with monolingual brain fine-tuning, it would suggest that the effects are driven by general brain representations rather than shared semantic representations in bilinguals. More details about datasets and preprocessing are in Appendix B.

### 3.2 Text-based Language Models

We fine-tuned monolingual and multilingual text-based Transformer language models to investigate the effects of brain-informed fine-tuning. All pretrained model checkpoints were obtained from Hugging Face (Wolf et al., 2020).

#### Monolingual pretrained language models

We fine-tuned two monolingual models: English BERT (BERT-en) and Chinese BERT (BERT-zh) (Devlin et al., 2019). Both models share the same architecture, comprising 12 Transformer layers with a hidden dimension size of 768, differing only in the language of their pretraining corpora.

#### Multilingual pretrained language models

We fine-tuned four multilingual Transformer-based models: mBERT (Devlin et al., 2019), XLM-R (Conneau et al., 2020), XGLM (Lin et al., 2022), and LLaMA-3.2 (Touvron et al., 2023). These models represent three distinct architecture types: encoder-based (mBERT, XLM-R), cross-lingual pretrained (XGLM), and decoder-based (LLaMA-3.2). Representations were extracted from the base versions of mBERT, XLM-R, and XGLM, and from the 1B parameter version of LLaMA-3.2. mBERT and XLM-R consist of 12 layers with a hidden dimension size of 768, XGLM has 24 layers with a hidden dimension size of 1024, and LLaMA-3.2 has 16 layers with a hidden dimension size of 2048. Results and analyses using XLM-R, XGLM and LLaMA-3.2 are reported in the Supplementary sections 2 and 3.

### 3.3 Brain-informed Fine-Tuning

We performed supervised full fine-tuning of pretrained language models using fMRI BOLD responses as targets. We employ a small-N design, where a language model is fine-tuned and evaluated independently for each participant, enabling robust within-participant inference (Smith & Little, 2018). An overview of the fine-tuning pipeline is shown in Fig. 1. We fine-tune language models using fMRI responses from either the whole-brain, language-selective (from Fedorenko et al. (2010)), or semantically-selective brain regions (see Appendix C Fig. 3). Details on fine-tuning with language-selective and semantically-selective brain regions are described in Appendix C.

**Figure 3:**
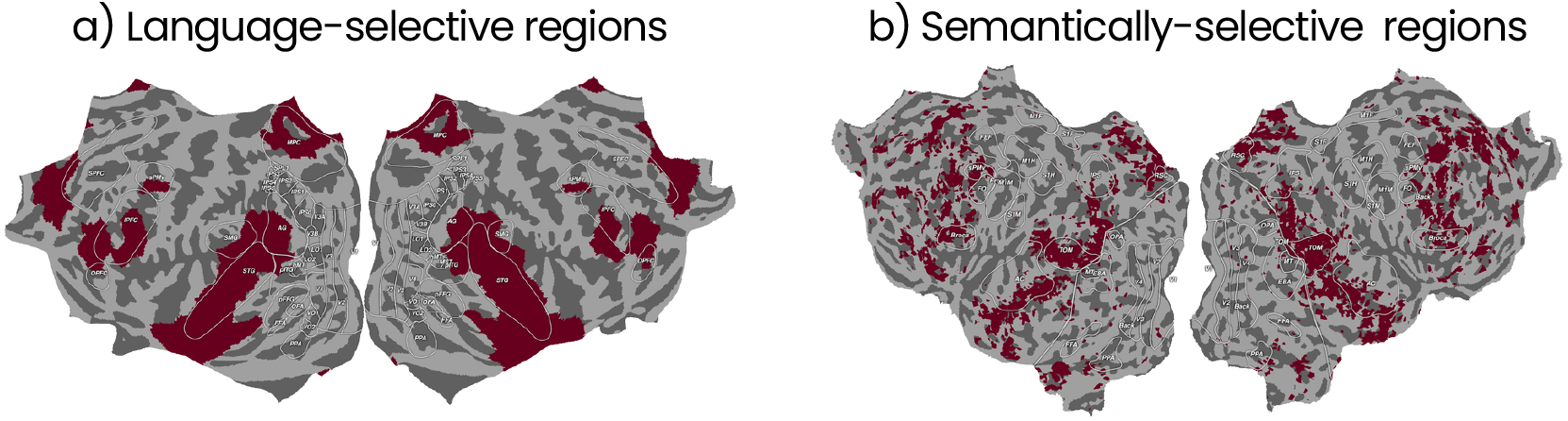
Cortical regions used for brain-informed fine-tuning. Flattened cortical surfaces for: (a) Language-selective regions displayed on the ‘fsaverage’ surface, used as the mask for all participants. and (b) semantically-selective regions for English for Participant 1. This mask is computed separately for each participant and language. Only fMRI responses from the dark red regions were used for fine-tuning in their respective variants.

**Figure 4:**
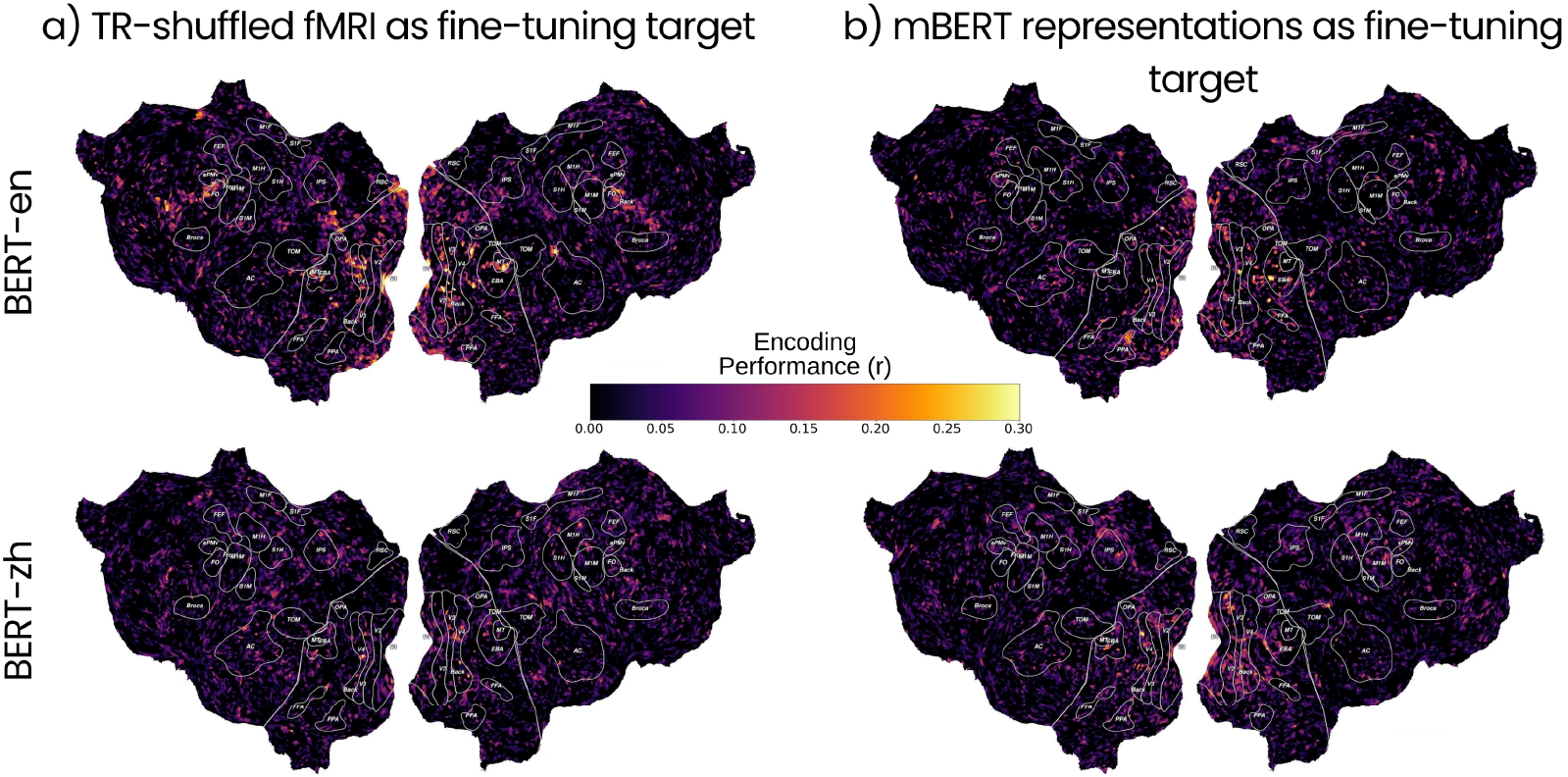
Encoding performance of baseline fine-tuned models. Voxelwise encoding performance is visualised for two fine-tuning baselines: (a) using TR-shuffled fMRI data, and (b) using mBERT representations as fine-tuning target. Flattened cortical surfaces with encoding performance are shown for Participant 1, with BERT-en (on top) and BERT-zh (on the bottom) fine-tuned with the baseline targets. In both baselines, the well predicted voxels do not correspond to semantically regions and appear scattered or in low-level sensory regions.

#### Model architecture

Text transcripts of the narrative stimuli were provided as inputs to the pretrained language model, using a sequence length of 20 tokens. We then extract the representations of the last token from the last hidden layer for each word. This representation is given as input to a dropout layer (a dropout of 0.2 is used to mitigate overfitting on limited brain data). This is followed by differentiable 3-lobe Lanczos (Huth et al., 2016) interpolation to downsample the text to the fMRI sampling rate. To model the temporal hemodynamic response, we learn a finite impulse response (FIR) filter by concatenating representations delayed by 2, 4, 6, and 8 seconds. The resulting representations are then passed through a linear projection layer to predict voxelwise BOLD responses. Previous brain-based fine-tuning studies (Moussa et al., 2025; Vattikonda et al., 2025) typically treated downsampling and temporal-delay modeling as separate preprocessing steps before model fine-tuning. In contrast, our method integrates these components directly into the model architecture, enabling end-to-end optimization. This design supports flexibility in handling variable-length inputs (as word timing per TR can vary) and allows for broader generalization across modalities and tasks. Comparison with a previously proposed fine-tuning pipeline is provided in the Supplementary section 4.

#### Training protocol

We fine-tuned the models using the AdamW optimizer (Loshchilov & Hutter, 2017) with a learning rate of 1e-4 and weight decay of 1e-3 for 30 epochs with a batch size of 32. A ReduceLROnPlateau scheduler was used to adjust the learning rate based on validation loss. We used mixed-precision training for computational efficiency and applied early stopping with a patience of 5 epochs based on validation performance. Implementation details are in Appendix D.

#### Training objective

Our training objective minimized the NT-Xent (Normalized Temperature-Scaled Cross-entropy) loss (Sohn, 2016) between the predicted and actual BOLD responses. We also experimented with alternative loss functions such as MSE, ridge loss, spatial loss, and a combination of all (referred to as hybrid), and found NT-Xent to perform best across models and experiments.

Results from other losses are reported in Supplementary section 1. We fine-tuned all layers of the language models and propagated the loss backward to update both the projection and transformer layers.

In addition to comparing our brain-informed fine-tuned models to their pretrained counterparts in both monolingual and multilingual settings, we include several baselines for a comprehensive comparison: TR-shuffled fMRI, multilingual model representations, and monolingual brain data as fine-tuning targets. This experiment was done to clarify the specific impact of bilingual brain data on the resulting fine-tuned model representations. Details about the baselines are reported in Appendix C.3.

### 3.4 Voxelwise Encoding Modeling

To evaluate whether brain-informed fine-tuning alters the alignment between language model representations and brain activity, we assess voxelwise encoding performance before and after fine-tuning. Voxelwise encoding models (VEM) estimate, for each voxel, a linear mapping from an embedding to the observed fMRI responses (Huth et al., 2016; Deniz et al., 2019). If fine-tuning with bilingual brain data improves encoding performance, we interpret this as evidence that the language model’s internal representations have become more brain-like, indicating successful alignment. If encoding performance remains unchanged, this suggests that fine-tuning preserves the existing brain alignment of the representations without introducing degradation. A decrease in performance would imply that fine-tuning introduces distortions that reduce the language model’s ability to predict brain responses, thus reducing its alignment with the brain.

We used representations extracted from layer 7 (found to yield the highest encoding performance based on validation data) as the embedding. Embeddings and brain responses were z-scored across time separately for each of the three stories not used for fine-tuning. Embeddings were downsampled to match the fMRI acquisition rate using 3-lobe Lanczos interpolation. Next, to account for the delayed hemodynamic response, each embedding was passed through a finite impulse response (FIR) filter with four delays. Specifically, delayed copies of each feature at 1, 2, 3, and 4 TRs (2, 4, 6, and 8 seconds) were concatenated. Ridge regression was used to determine how the embedding is represented in each voxel (Wu et al., 2006; Naselaris et al., 2011). We performed 5-fold cross-validation to find optimal regularization parameters. Encoding performance was quantified by calculating the Pearson correlation coefficient (r) between predicted and recorded BOLD responses using a held-out story. A separate VEM was fit for each voxel, participant, and language model variant (before and after brain-informed fine-tuning). No sample size calculations were performed, as each participant serves as a full replication of the results.

#### Cross-participant transfer of encoding performance

We evaluated whether brain-informed fine-tuned model representations generalize across participants. We fine-tuned the language model using brain data from one bilingual participant and evaluated its encoding performance using VEMs on other participants. Improved transfer would suggest that changes introduced by brain-informed fine-tuning are not participant-specific but reflect shared representations across bilingual individuals.

### 3.5 Downstream NLP tasks

To evaluate how brain-informed fine-tuning changes language model representations, we assessed its performance on standard NLP benchmarks before and after fine-tuning. For English, we used 9 tasks from the GLUE benchmark (Wang et al., 2018), and for Chinese, we used 7 tasks from the CLUE benchmark (Xu et al., 2020). Both monolingual and multilingual models were evaluated on these benchmarks. To investigate cross-linguistic generalization to unseen languages, we additionally evaluated multilingual models on 3 tasks from the XGLUE benchmark (Liang et al., 2020) and 3 tasks from the XTREME benchmark (Hu et al., 2020). Details about these benchmarks and task metrics are in Appendix E.

#### Fine-tuning and evaluation in the same language

To assess how brain-informed fine-tuning of language models affects performance on NLP tasks within the same language, we fine-tuned language models using fMRI data in English or Chinese. We then evaluated these language models on NLP benchmarks in the corresponding language. English and Chinese brain data are used to fine-tune monolingual (BERT-en, BERT-zh) and multilingual (mBERT) language models, which are then evaluated on GLUE and CLUE, respectively. Any performance increase observed in this setting would imply that brain-informed fine-tuning not only changes language model representations to better align with the brain, but that these modified representations are better for NLP tasks within the trained language.

#### Cross-language transfer between known languages

To test whether brain-informed fine-tuning transfers to a bilingual participant’s other language, we evaluate models on the participant’s other language after fine-tuning on the first. For example, weights from BERT-en fine-tuned with English brain data are transferred to BERT-zh and evaluated on Chinese tasks, and vice versa. Improved performance in this setting would suggest that brain-informed fine-tuning introduces the bilingual brain’s shared semantic representations into the model, enabling cross-language transfer.

#### Zero-shot transfer to unseen languages

To evaluate the broader generalizability of brain-informed fine-tuned models, we tested multilingual models on five additional languages (German, French, Spanish, Japanese, and Korean) that were neither used for fine-tuning nor known to the participant. Specifically, we fine-tuned mBERT on English brain data and evaluated its performance on the XGLUE (Liang et al., 2020) and XTREME (Hu et al., 2020) benchmarks. This setup allows us to assess whether brain-informed fine-tuning can introduce language-agnostic representations that support broader cross-linguistic transfer.

## 4 Results

### 4.1 Effects on Brain Encoding Performance and Generalization

We tested whether brain-informed fine-tuning improves brain encoding performance and if these possible improvements are generalizable across participants. We compared the VEM performance of the vanilla language model against the three fine-tuned variants, trained using whole-brain or language-selective, or participant-specific semantically-selective voxels. Fig. 2 shows flattened cortical surfaces for participant 1, visualizing which language model variant achieved the best encoding performance across well-predicted voxels (Pearson correlation *r*>0.1) for (a) monolingual and (b) multilingual models within the same language. Across both languages and model types, we found that fine-tuned language models outperform their vanilla counterparts, with the best-performing variant explaining more variance in a majority of voxels (76-84% of well-predicted voxels). This was consistent across all six participants (see Fig. 5 in Appendix F.2 for VEM results on other participants). Our results suggest that a language model’s ability to predict brain activity benefits from brain-informed fine-tuning. Further, as shown in Appendix F.1, brain-informed fine-tuning outperforms control baselines (like TR-shuffled fMRI responses and mBERT representations as fine-tuning targets).

**Figure 5:**
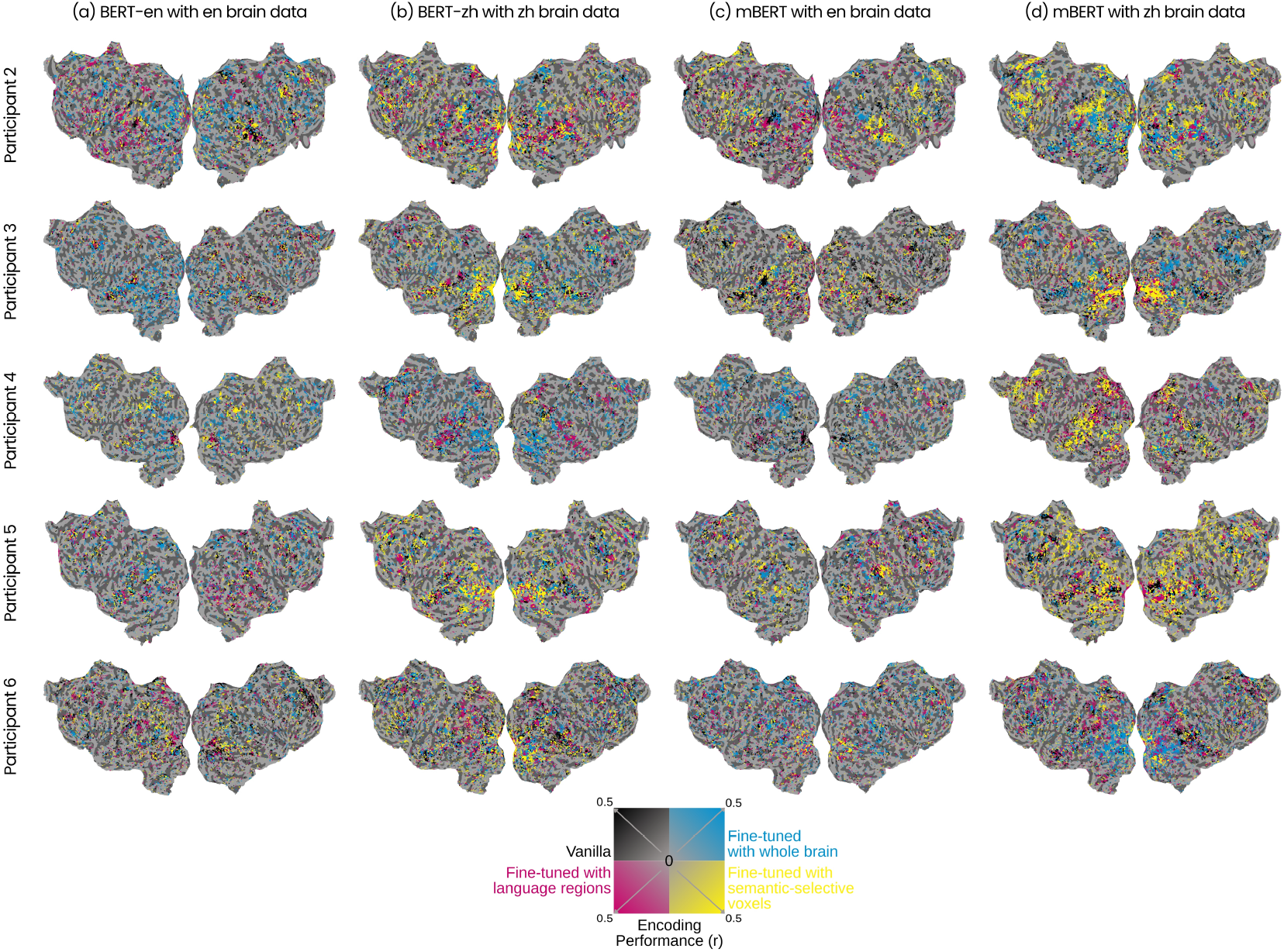
Brain encoding performance before and after brain-informed fine-tuning for all participants. This figure shows voxelwise encoding model performance for the best-performing fine-tuned variants across the cortex for all participants, using (a) BERT-en, (b) BERT-zh, and (c–d) mBERT models. This replicates the format of Fig. 2, and reports it for all participants. For details on voxel color coding, refer to the caption of Fig. 2. Across participants, model types, and languages, brain-informed fine-tuning consistently improves encoding performance relative to it’s vanilla counterpart.

We do not observe any regional preference for any particular fine-tuning approach. No specific brain region consistently preferred one fine-tuned variant over the others across participants. Although fine-tuning improved encoding performance, the overall increase was modest (maximum Δ*r* ≈ 0.15), which aligns with prior studies. For example, previous work has shown that current representations from language models already serve as one of the best predictors of brain responses (Schrimpf et al., 2021), and even in speech models that lack brain-relevant semantics (Oota et al., 2024a), fine-tuning yielded small improvements (Δ*r ≈* 0.1) (Vattikonda et al., 2025).

To test whether these improvements were participant-specific, we checked for cross-participant generalization. We fine-tuned a BERT-en model on one participant’s English brain data and evaluated its VEM performance on the other participants. As shown in Fig. 6 in Appendix F.3, we observe small but consistent improvements (Δ*r* ≈ 0.03 −0.05) in high-level semantic areas. These findings suggest that the improvements from brain-informed fine-tuning of language model representations are not specific to individual participants but reflect shared representations across bilingual individuals.

**Figure 6:**
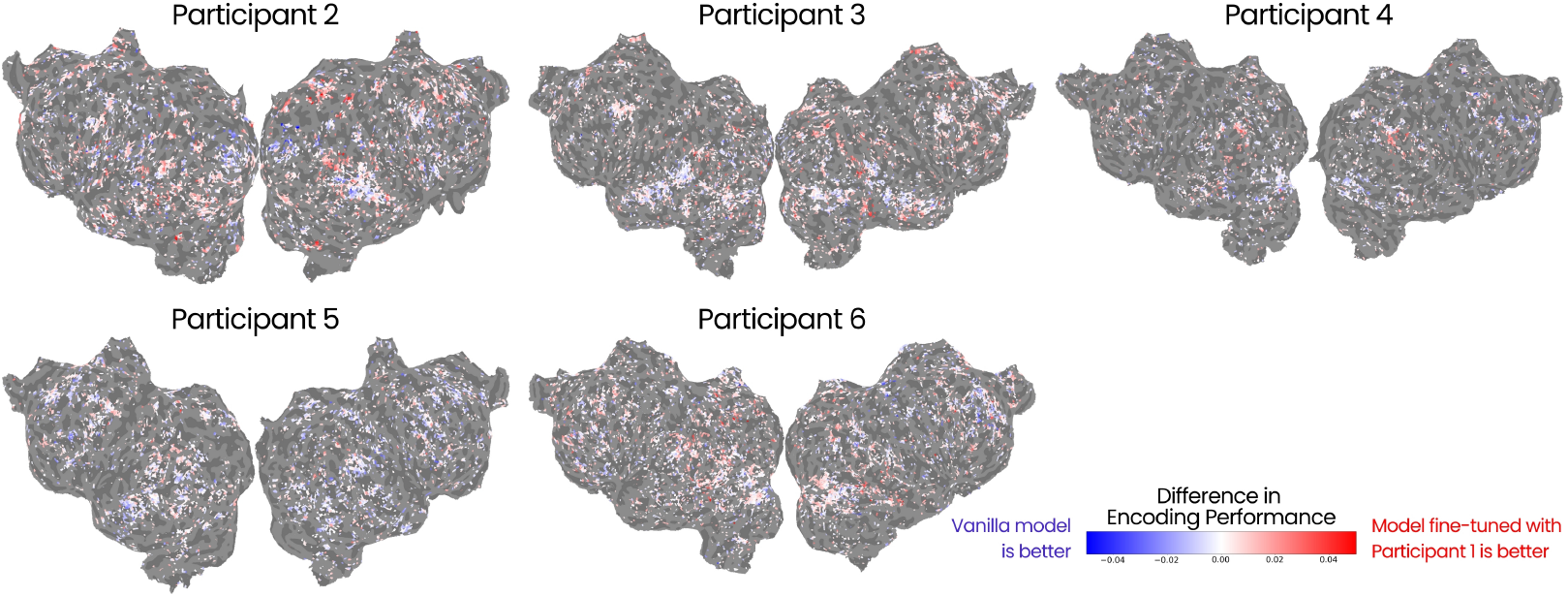
Cross-participant generalization of brain-informed fine-tuned language models. Flattened cortical surfaces show changes in encoding performance for Participants 2–6 when comparing BERT-en (vanilla) with BERT-ft-en fine-tuned using English brain data from Participant 1. Each point represents a voxel that was consistently well-predicted (r*>*0.1) across both models. Across participants, encoding performance in higher-level semantic regions shows either no degradation or slight improvements with the fine-tuned model.

### 4.2 Performance on Downstream NLP tasks

We tested the effect of brain-informed fine-tuning (with semantically-selective brain regions) on language models by evaluating model performance on standard NLP benchmarks. We assess this in three settings:

#### Fine-tuning and evaluation in the same language

We show in Table 1(a) performance on down-stream NLP tasks when brain-informed fine-tuning (with semantically-selective brain regions) and evaluation are computed in the same language. For English, models were fine-tuned using English brain data and evaluated on the GLUE benchmark. We observe that the monolingual model (BERT-ft-en) outperforms its vanilla counterpart (BERT-en) on 7/9 tasks, with an average improvement of +0.80 percentage points, and a maximum gain of +3.57 on the WNLI task. The multilingual model (mBERT-ft-en) improves on 7/9 tasks, with an average gain of +0.89 percentage points, and max gain of +3.12 also on WNLI. For Chinese, models were fine-tuned using Chinese brain data and evaluated on the CLUE benchmark. Here, BERT-ft-zh achieves slight improvements on 5/7 tasks relative to BERT-zh, with an average gain of +0.65 percentage point and max gain of +1.23 on the CSL task. The multilingual model (mBERT-ft-zh) also improves on 6/7 tasks, achieving an average improvement of +0.53 percentage point, with the largest improvement of +1.02 observed on the C^3^ task. These results indicate that brain-informed fine-tuning consistently improves downstream task performance for both monolingual and multilingual models when the fine-tuning and evaluation languages match.

**Table 1:** Downstream task performance before and after bilingual brain-informed fine-tuning (with semantically-selective brain regions). We compared vanilla (pretrained) models, English BERT (BERT-en), Chinese BERT (BERT-zh), and mBERT, with their bilingual brain-informed fine-tuned counterparts (e.g., BERT-ft-en: BERT-en fine-tuned using English brain data from semantically-selective brain regions). For each task, the average performance and standard deviation across the six bilingual participants are reported. To assess whether brain-informed fine-tuning elicits multilingual capabilities, we evaluate downstream task performance in three settings: (a) Fine-tuning and evaluation in the same language: models are fine-tuned with brain data in one language (en or zh) and evaluated on NLP tasks in the same language (GLUE benchmark for en, CLUE benchmark for zh). This tests within-language improvements due to brain-informed fine-tuning. (b) Cross-language transfer between known languages: model is fine-tuned on brain data in one language and evaluated on tasks in the participants’ second (not used in fine-tuning) language (e.g., fine-tuned with en brain data and evaluated on CLUE (zh benchmark) tasks). This tests whether bilingual brain-informed fine-tuning elicits the participants’ shared semantic representations. (c) Zero-shot transfer to unseen languages: to assess broader multilingual transfer, mBERT-ft-en is evaluated on downstream tasks in additional languages not seen during fine-tuning (German, French, Spanish, Japanese, and Korean) using XGLUE and XTREME benchmarks. Bolded values indicate performance equal to or better than the corresponding vanilla model. We observe that bilingual brain-informed fine-tuning improves performance on several NLP tasks, with mBERT showing greater benefits than BERT across all three settings. Results for fine-tuning using whole-brain and language-selective regions are reported in Appendix G.1; see Tables 9 and 10.

**a) Fine-tuning and Evaluation in the Same Language**
| GLUE Task | BERT-en | BERT-ft-en | mBERT | mBERT-ft-en |
| --- | --- | --- | --- | --- |
| CoLA (MCC) | 53.38 | **55.25 ± 0.86** | **42.68** | 41.89 ± 0.74 |
| SST-2 (Acc.) | 92.08 | **92.62 ± 0.10** | 89.68 | **91.06 ± 0.44** |
| MRPC (Acc.) | 79.41 | **81.43 ± 0.79** | 84.80 | **85.96 ± 0.59** |
| MRPC (F1) | 86.27 | **87.47 ± 0.49** | 88.56 | **89.22 ± 0.66** |
| STS-B (Pears.) | **88.06** | 87.70 ± 0.19 | 88.06 | **88.31 ± 0.77** |
| STS-B (Spear.) | **87.65** | 87.54 ± 0.20 | 87.76 | **88.27 ± 0.69** |
| QQP (Acc.) | 90.84 | **90.88 ± 0.03** | 90.22 | **90.47 ± 0.04** |
| QQP (F1) | 87.70 | **87.79 ± 0.02** | 86.70 | **87.15 ± 0.07** |
| MNLI-m (Acc.) | **84.38** | 84.29 ± 0.26 | 82.09 | **82.10 ± 0.19** |
| MNLI-mm (Acc.) | **84.64** | 84.60 ± 0.31 | 82.38 | **82.52 ± 0.34** |
| QNLI (Acc.) | 91.45 | **91.53 ± 0.06** | 91.14 | **91.18 ± 0.16** |
| RTE (Acc.) | 67.15 | **67.47 ± 0.46** | **67.15** | 66.78 ± 1.15 |
| WNLI (Acc.) | 49.30 | **52.87 ± 0.64** | 53.52 | **56.64 ± 0.62** |

| CLUE Task | BERT-zh | BERT-ft-zh | mBERT | mBERT-ft-zh |
| --- | --- | --- | --- | --- |
| AFQMC (Acc.) | **75.25** | 73.76 ± 0.39 | 69.74 | **70.37 ± 0.38** |
| CMNLI (Acc.) | 80.50 | **80.93 ± 0.20** | 78.66 | **79.22 ± 0.25** |
| CSL (F1) | 80.18 | **81.41 ± 0.36** | 81.10 | **81.36 ± 0.59** |
| IFLYTEK (Acc.) | 60.25 | **60.37 ± 0.50** | 56.52 | **56.66 ± 0.19** |
| TNEWS (Acc.) | 56.24 | **56.50 ± 0.14** | 54.77 | **55.02 ± 0.17** |
| TNEWS (F1) | 55.17 | **55.95 ± 0.27** | 53.69 | **54.07 ± 0.24** |
| ChID (Acc.) | 10.66 | **10.66 ± 0.00** | 10.66 | **10.66 ± 0.00** |
| C<sup>3</sup> (Acc.) | 49.74 | **49.79 ± 0.42** | 49.42 | **50.44 ± 0.96** |

**b) Cross-Language Transfer Between Known Languages**
| GLUE Task | BERT-zh | BERT-ft-zh | mBERT | mBERT-ft-zh |
| --- | --- | --- | --- | --- |
| CoLA (MCC) | 51.13 | **52.73 ± 0.56** | 40.96 | **41.18 ± 0.41** |
| SST-2 (Acc.) | 92.26 | **92.59 ± 0.13** | 88.27 | **90.51 ± 0.33** |
| MRPC (Acc.) | 76.96 | **78.90 ± 0.45** | 82.84 | **85.80 ± 0.23** |
| MRPC (F1) | 83.38 | **85.08 ± 0.71** | 87.15 | **89.45 ± 0.47** |
| STS-B (Pears.) | 87.21 | **88.37 ± 0.18** | 87.09 | **88.09 ± 0.44** |
| STS-B (Spear.) | 86.73 | **88.16 ± 0.13** | 86.97 | **87.99 ± 0.48** |
| QQP (Acc.) | 90.65 | **91.03 ± 0.36** | 89.91 | **90.45 ± 0.08** |
| QQP (F1) | 87.53 | **87.92 ± 0.20** | 86.27 | **87.08 ± 0.10** |
| MNLI-m (Acc.) | 84.00 | **84.59 ± 0.42** | 81.52 | **82.07 ± 0.09** |
| MNLI-mm (Acc.) | 83.91 | **84.25 ± 0.46** | 81.94 | **82.74 ± 0.12** |
| QNLI (Acc.) | 91.27 | **91.35 ± 0.05** | 90.55 | **91.04 ± 0.10** |
| RTE (Acc.) | **67.15** | 66.79 ± 0.65 | 66.06 | **68.71 ± 0.95** |
| WNLI (Acc.) | 53.52 | **56.34 ± 0.00** | 54.93 | **56.34 ± 0.00** |

| CLUE Task | BERT-en | BERT-ft-en | mBERT | mBERT-ft-en |
| --- | --- | --- | --- | --- |
| AFQMC (Acc.) | 69.00 | **69.31 ± 0.38** | 69.74 | **71.12 ± 0.54** |
| CMNLI (Acc.) | 68.34 | **68.63 ± 0.21** | 78.66 | **79.09 ± 0.14** |
| CSL (F1) | 71.20 | **71.63 ± 1.55** | 81.10 | **81.34 ± 0.49** |
| IFLYTEK (Acc.) | **47.86** | 47.53 ± 0.27 | 56.52 | **57.21 ± 0.51** |
| TNEWS (Acc.) | 50.92 | **51.48 ± 0.49** | 54.77 | **54.93 ± 0.72** |
| TNEWS (F1) | 50.15 | **50.78 ± 0.92** | 53.69 | **53.97 ± 0.26** |
| ChID (Acc.) | 10.66 | **10.66 ± 0.00** | 10.66 | **10.66 ± 0.00** |
| C<sup>3</sup> (Acc.) | **42.64** | 41.05 ± 6.41 | 49.42 | **50.33 ± 0.43** |

**c) Zero-Shot Transfer to Unseen Languages**
| Task | German (de) |  | French (fr) |  | Spanish (es) |  | Japanese (ja) |  | Korean (ko) |  |
| --- | --- | --- | --- | --- | --- | --- | --- | --- | --- | --- |
|  | mBERT | mBERT-ft-en | mBERT | mBERT-ft-en | mBERT | mBERT-ft-en | mBERT | mBERT-ft-en | mBERT | mBERT-ft-en |
| **XGLUE** |  |  |  |  |  |  |  |  |  |  |
| PAWS-X (Acc.) | 84.00 | **84.90 ± 0.54** | 88.10 | **88.48 ± 0.44** | 87.05 | **87.44 ± 0.48** | 78.70 | **78.70 ± 0.17** | **78.75** | 78.36 ± 1.43 |
| XNLI (Acc.) | 70.60 | **71.04 ± 0.83** | 72.69 | **73.38 ± 0.91** | 74.10 | **74.96 ± 0.54** | 86.00 | **88.53 ± 0.15** | 86.00 | **88.25 ± 0.12** |
| NER (Acc.) | 10.67 | **11.89 ± 0.77** | 06.67 | **11.78 ± 1.68** | 07.33 | **12.40 ± 1.82** | **13.00** | 10.89 ± 0.51 | **12.00** | 11.13 ± 1.48 |
| **XTREME** |  |  |  |  |  |  |  |  |  |  |
| PAWS-X (Acc.) | **85.65** | 84.51 ± 0.01 | **88.95** | 88.25 ± 0.01 | **87.90** | 87.18 ± 0.01 | **79.55** | 78.98 ± 0.01 | **79.35** | 78.71 ± 0.01 |
| XNLI (Acc.) | 70.06 | **71.32 ± 0.86** | 72.97 | **73.53 ± 0.25** | 74.50 | **74.73 ± 0.13** | 86.00 | **87.13 ± 0.10** | 86.00 | **87.21 ± 0.51** |
| NER (Acc.) | 12.33 | **12.94 ± 0.83** | 09.33 | **11.91 ± 1.03** | 10.33 | **11.89 ± 0.46** | 08.67 | **11.09 ± 0.92** | **12.33** | 11.87 ± 0.77 |

**Table 2:**
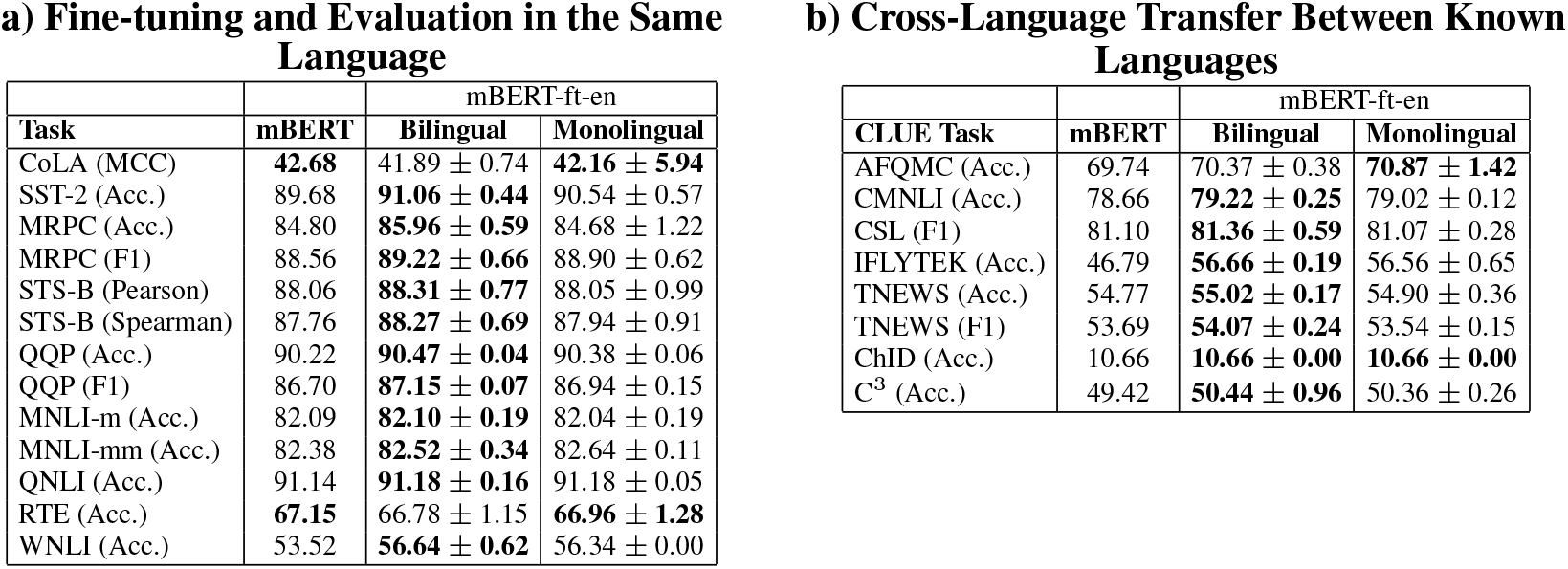
Downstream task performance before and after bilingual or monolingual brain-informed fine-tuning (with semantically-selective brain regions). We perform brain-informed fine-tuning of mBERT with English brain data (mBERT-ft-en) from semantically-selective brain regions, from either bilingual or monolingual participants. For each task, the average performance and standard deviation across the participants are reported. We evaluate downstream task performance in two settings: (a) Fine-tuning and evaluation in the same language: the model is evaluated in English with GLUE tasks. (b) Cross-language transfer between known languages: the model is evaluated in Chinese with CLUE tasks.

**Table 3:** Descriptions of tasks in the GLUE and CLUE benchmarks.

| Task | Description | Task Type |
| --- | --- | --- |
| **GLUE Benchmark** |  |  |
| CoLA | Determine linguistic acceptability of a sentence | Single-sentence classification |
| SST-2 | Sentiment analysis on movie reviews | Single-sentence classification |
| MRPC | Determine if two sentences are semantically equivalent | Sentence-pair classification |
| STS-B | Measure semantic similarity (0–5) between two sentences | Sentence-pair regression |
| QQP | Identify if two questions are duplicates | Sentence-pair classification |
| MNLI | Determine if a premise entails, contradicts, or is neutral to a hypothesis | Sentence-pair classification (multi-class) |
| QNLI | Determine if a sentence contains the answer to a question | Sentence-pair classification |
| RTE | Determine textual entailment between two sentences | Sentence-pair classification (binary) |
| WNLI | Coreference resolution: determine if a hypothesis is true given a sentence with a replaced pronoun | Sentence-pair classification |
| **CLUE Benchmark** |  |  |
| AFQMC | Determine if two Chinese sentences are semantically equivalent | Sentence-pair classification |
| CMNLI | Multilingual version of MNLI for Chinese | Sentence-pair classification |
| CSL | Determine whether a keyword list matches a given abstract | Sentence-pair classification |
| IFLYTEK | Classify app descriptions into predefined categories | Multi-class classification |
| TNEWS | Classify news headlines into categories | Multi-class classification |
| ChID | Fill-in-the-blank multiple-choice cloze test from Chinese literature | Cloze-style multiple-choice |
| C<sup>3</sup> | Reading comprehension with multiple-choice answers | Multi-turn QA / Multiple-choice |

**Table 4:** Summary of XGLUE and XTREME tasks used in this study.

| Benchmark | Task Name | Task Type | Languages Evaluated | Evaluation Metric |
| --- | --- | --- | --- | --- |
| XGLUE | PAWS-X | Paraphrase Identification | de, fr, es, ja, ko | Accuracy |
|  | XNLI | Natural Language Inference | de, fr, es, ja, ko | Accuracy |
|  | NER | Named Entity Recognition | de, fr, es, ja, ko | F1 Score |
| XTREME | PAWS-X | Paraphrase Identification | de, fr, es, ja, ko | Accuracy |
|  | XNLI | Natural Language Inference | de, fr, es, ja, ko | Accuracy |
|  | NER | Named Entity Recognition | de, fr, es, ja, ko | F1 Score |

**Table 5:** GLUE tasks, metrics, reference ranges, and scaling.

| GLUE Task | Metric | Reference Range | Scaled |
| --- | --- | --- | --- |
| CoLA | Matthews Correlation | $[-1, 1]$ | $\times 100$ |
| SST-2 | Accuracy | $[0, 1]$ | $\times 100$ |
| MRPC | F1 Score / Accuracy | $[0, 1]$ | $\times 100$ |
| STS-B | Pearson / Spearman Correlation | $[-1, 1]$ | $\times 100$ |
| QQP | F1 Score / Accuracy | $[0, 1]$ | $\times 100$ |
| MNLI | Accuracy | $[0, 1]$ | $\times 100$ |
| QNLI | Accuracy | $[0, 1]$ | $\times 100$ |
| RTE | Accuracy | $[0, 1]$ | $\times 100$ |
| WNLI | Accuracy | $[0, 1]$ | $\times 100$ |

**Table 6:** CLUE tasks, metrics, reference ranges, and scaling.

| CLUE Task | Metric | Reference Range | Scaled |
| --- | --- | --- | --- |
| AFQMC | Accuracy | $[0, 1]$ | $\times 100$ |
| CMNLI | Accuracy | $[0, 1]$ | $\times 100$ |
| CSL | F1 Score | $[0, 1]$ | $\times 100$ |
| IFLYTEK | Accuracy | $[0, 1]$ | $\times 100$ |
| TNEWS | Accuracy / F1 Score | $[0, 1]$ | $\times 100$ |
| ChID | Accuracy | $[0, 1]$ | $\times 100$ |
| C<sup>3</sup> | Exact Match / Accuracy | $[0, 1]$ | $\times 100$ |

**Table 7:** XGLUE tasks, metrics, reference ranges, and scaling.

| XGLUE Task | Metric | Reference Range | Scaled |
| --- | --- | --- | --- |
| NER | F1 Score | $[0, 1]$ | $\times 100$ |
| PAWS-X | Accuracy | $[0, 1]$ | $\times 100$ |
| XNLI | Accuracy | $[0, 1]$ | $\times 100$ |

#### Cross-language transfer between known languages

Table 1(b) reports performance when brain-informed fine-tuning (with semantically-selective brain regions) is done in one language and evaluation is performed in the participant’s other language. On the GLUE benchmark, we evaluated BERT-zh and mBERT models fine-tuned using Chinese brain data. We observe that the monolingual model (BERT-ft-zh) outperforms its vanilla counterpart (BERT-zh) on 8/9 tasks, with an average improvement of +1.01 percentage points, and a maximum gain of +2.82 on the WNLI task. The multilingual model (mBERT-ft-zh) improves on all tasks, with an average gain of +1.61 percentage points, and max gain of +2.96 on MRPC (Acc.). On the CLUE benchmark, we evaluated BERT-en and mBERT models fine-tuned using English brain data. BERT-ft-en shows improvement on 4/7 tasks, with a mean improvement of +0.44 points overall (max gain: +0.63 on TNEWS (F1)). mBERT-ft-en improves across 6/7 tasks, with an average gain of +0.68 points and a maximum improvement of +1.38 on the AFQMC task.

Overall, bilingual brain-informed fine-tuning enables cross-lingual generalization in language models, improving downstream task performance even when the fine-tuning and evaluation languages differ. This suggests that bilingual brain-informed fine-tuning of language models introduces changes that reflect the bilingual brain’s shared semantic representations, enabling cross-language transfer.

#### Zero-shot transfer to unseen languages

Table 1(c) reports performance on downstream NLP tasks when brain-informed fine-tuning (with semantically-selective brain regions) is performed with English brain data and evaluated on five unseen languages (German, French, Spanish, Japanese, and Korean), that were neither used for fine-tuning nor known to the participant. We evaluate the multilingual model (mBERT-ft-en) using tasks from the XGLUE and XTREME benchmarks. On the XGLUE benchmark, mBERT-ft-en improves performance on 3/3 tasks in German, French, and Spanish, and 1/3 tasks in Japanese and Korean, with average gains of +0.85, +2.06, +2.11, +0.14, and +0.33 percentage points, respectively. On the XTREME benchmark, mBERT-ft-en improves performance on 2/3 tasks in German, French, Spanish, and Japanese and 1/3 tasks in Korean, with average gains of +0.24, +0.81, +0.36, +0.99, and +0.04 percentage points, respectively. These results demonstrate that bilingual brain-informed fine-tuning improves language-agnostic representations in multilingual models, enabling generalization beyond the language of the brain data to entirely unseen languages (zero-shot).

#### Performance on downstream tasks with whole-brain or language-selective voxels

We evaluated downstream task performance before and after brain-informed fine-tuning using either whole-brain or language-selective voxels. Results averaged across all participants (mean ± std) for these two settings are reported in Appendix G.1, Tables 9 and 10, respectively. Table 8 in Appendix G further compares mBERT-en (fine-tuned with English brain data) and mBERT-zh (fine-tuned with Chinese brain data) when using either whole-brain, language-selective, or semantically-selective regions for fine-tuning. Fine-tuning with either variant improves performance across all downstream tasks compared to vanilla models.

**Table 8:** Downstream task performance before and after bilingual brain-informed fine-tuning (with whole-brain, or language-selective, or semantically-selective regions).

a) Fine-tuning and Evaluation in the Same Language
| Task | mBERT-en | Whole-Brain | Language | Semantic | CLUE Task | mBERT-zh | Whole-Brain | Language | Semantic |
| --- | --- | --- | --- | --- | --- | --- | --- | --- | --- |
| CoLA (MCC) | 42.68 | 40.05 | 41.80 | **42.81** | AFQMC (Acc.) | 69.74 | 70.55 | **72.03** | 71.11 |
| SST-2 (Acc.) | 89.68 | 90.14 | 90.25 | **90.60** | CMNLI (Acc.) | 78.66 | **79.02** | 78.75 | 79.01 |
| MRPC (Acc.) | 84.80 | 84.80 | **86.76** | 86.03 | CSL (F1) | 81.10 | 80.71 | **82.27** | 81.24 |
| MRPC (F1) | 88.56 | 89.01 | **90.49** | 89.73 | IFLYTEK (Acc.) | 56.52 | 56.83 | 56.79 | **56.91** |
| STS-B (Pearson) | 88.06 | 88.42 | **88.46** | 88.42 | TNEWS (Acc.) | 54.77 | 55.12 | **55.27** | 54.91 |
| STS-B (Spearman) | 87.76 | 88.22 | **88.38** | 88.34 | TNEWS (F1) | 53.69 | 53.74 | 53.82 | **53.98** |
| QQP (Acc.) | 90.22 | 90.47 | 90.43 | **90.48** | ChID (Acc.) | 10.66 | **10.66** | **10.66** | **10.66** |
| QQP (F1) | 86.70 | 87.07 | 87.02 | **87.11** | C<sup>3</sup> (Acc.) | 49.42 | **50.21** | 49.76 | 49.95 |
| MNLI-m (Acc.) | 82.09 | **82.66** | 82.27 | 82.07 |  |  |  |  |  |
| MNLI-mm (Acc.) | 82.38 | **82.97** | 82.92 | 82.68 |  |  |  |  |  |
| QNLI (Acc.) | 91.14 | 91.18 | 91.10 | **91.31** |  |  |  |  |  |
| WNLI (Acc.) | 53.52 | **56.34** | **56.34** | **56.34** |  |  |  |  |  |
| RTE (Acc.) | 67.15 | 65.70 | **68.95** | 67.87 |  |  |  |  |  |

(b) Cross-Language Transfer Between Known Languages
| Task | mBERT-zh | Whole-Brain | Language | Semantic | CLUE Task | mBERT-en | Whole-Brain | Language | Semantic |
| --- | --- | --- | --- | --- | --- | --- | --- | --- | --- |
| CoLA (MCC) | 40.96 | 41.22 | 40.82 | **41.60** | AFQMC (Acc.) | 69.74 | 70.62 | **71.29** | 70.76 |
| SST-2 (Acc.) | 88.27 | 89.25 | **91.17** | 90.37 | CMNLI (Acc.) | 78.66 | 78.83 | 78.92 | **79.09** |
| MRPC (Acc.) | 82.84 | 83.82 | 85.05 | **87.01** | CSL (F1) | 81.10 | 81.30 | 81.53 | **81.57** |
| MRPC (F1) | 87.15 | 87.88 | 89.09 | **90.62** | IFLYTEK (Acc.) | 46.79 | 56.52 | **56.60** | 56.33 |
| STS-B (Pearson) | 87.09 | 88.11 | 87.00 | **88.54** | TNEWS (Acc.) | 54.77 | 54.81 | **54.95** | 53.47 |
| STS-B (Spearman) | 86.97 | 87.83 | 87.27 | **88.43** | TNEWS (F1) | 53.69 | 53.84 | **54.01** | 53.75 |
| QQP (Acc.) | 89.91 | 90.16 | **90.46** | 90.42 | ChID (Acc.) | 10.66 | **10.66** | **10.66** | **10.66** |
| QQP (F1) | 86.27 | 86.89 | **87.06** | 87.00 | C<sup>3</sup> (Acc.) | 49.42 | 49.48 | **49.84** | 49.76 |
| MNLI-m (Acc.) | 81.52 | 82.13 | 82.54 | **82.94** |  |  |  |  |  |
| MNLI-mm (Acc.) | 81.94 | 82.36 | 82.74 | **83.12** |  |  |  |  |  |
| QNLI (Acc.) | 90.55 | 90.92 | 91.27 | **91.38** |  |  |  |  |  |
| WNLI (Acc.) | 54.93 | **56.34** | **56.34** | **56.34** |  |  |  |  |  |
| RTE (Acc.) | 66.06 | 66.43 | **68.59** | 66.06 |  |  |  |  |  |

**Table 9:** Downstream task performance before and after bilingual brain-informed fine-tuning (with whole-brain). We compared vanilla (pretrained) models, English BERT (BERT-en), Chinese BERT (BERT-zh), and mBERT, with their bilingual brain-informed fine-tuned counterparts (e.g., BERT-ft-en: BERT-en fine-tuned using English brain data). For each task, the average performance and standard deviation across the six bilingual participants are reported. To assess whether brain-informed fine-tuning elicits multilingual capabilities, we evaluate downstream task performance in three settings: (a) Fine-tuning and evaluation in the same language: models are fine-tuned with brain data in one language (en or zh) and evaluated on NLP tasks in the same language (GLUE benchmark for en, CLUE benchmark for zh). This tests within-language improvements due to brain-informed fine-tuning. (b) Cross-language transfer between known languages: model is fine-tuned on brain data in one language and evaluated on tasks in the participants’s second (not used in fine-tuning) language (e.g., fine-tuned with en brain data and evaluated on CLUE (zh benchmark) tasks). This tests whether bilingual brain-informed fine-tuning elicits the participants’ shared semantic representations. (c) Zero-shot transfer to unseen languages: to assess broader multilingual transfer, mBERT-ft-en is evaluated on downstream tasks in additional languages not seen during fine-tuning (German, French, Spanish, Japanese, and Korean) using XGLUE and XTREME benchmarks. Bolded values indicate performance equal to or better than the corresponding vanilla model. We observe that bilingual brain-informed fine-tuning improves performance on several NLP tasks, with mBERT showing greater benefits than BERT across all three settings. Results for fine-tuning using whole-brain data for individual participants are reported in the Supplementary (see Table 4).

**a) Fine-tuning and Evaluation in the Same Language**
| GLUE Task | BERT-en | BERT-ft-en | mBERT | mBERT-ft-en |
| --- | --- | --- | --- | --- |
| CoLA (MCC) | 53.38 | **55.11 ± 0.75** | **42.68** | 39.58 ± 1.82 |
| SST-2 (Acc.) | 92.08 | **92.58 ± 0.39** | 89.68 | **90.64 ± 0.54** |
| MRPC (Acc.) | 79.41 | **80.55 ± 1.01** | 84.80 | **85.39 ± 0.51** |
| MRPC (F1) | 86.27 | **86.91 ± 0.75** | 88.56 | **89.20 ± 0.27** |
| STS-B (Pears.) | 88.06 | **88.10 ± 0.07** | 88.06 | **88.47 ± 0.09** |
| STS-B (Spear.) | **87.65** | 87.55 ± 0.18 | 87.76 | **88.28 ± 0.12** |
| QQP (Acc.) | **90.84** | 90.79 ± 0.03 | 90.22 | **90.43 ± 0.04** |
| QQP (F1) | 87.70 | **87.71 ± 0.19** | 86.70 | **87.08 ± 0.02** |
| MNLI-m (Acc.) | 84.38 | **84.40 ± 0.09** | 82.09 | **82.26 ± 0.27** |
| MNLI-mm (Acc.) | **84.64** | 84.55 ± 0.20 | 82.38 | **82.73 ± 0.14** |
| QNLI (Acc.) | 91.45 | **91.49 ± 0.09** | 91.14 | **91.24 ± 0.06** |
| RTE (Acc.) | 67.15 | **67.32 ± 0.17** | **67.15** | 65.63 ± 0.51 |
| WNLI (Acc.) | 49.30 | **55.59 ± 1.31** | 53.52 | **56.34 ± 0.00** |

| CLUE Task | BERT-zh | BERT-ft-zh | mBERT | mBERT-ft-zh |
| --- | --- | --- | --- | --- |
| AFQMC (Acc.) | **75.25** | 74.50 ± 0.48 | 69.74 | **70.70 ± 0.99** |
| CMNLI (Acc.) | 80.50 | **80.87 ± 0.04** | 78.66 | **78.99 ± 0.14** |
| CSL (F1) | 80.18 | **80.82 ± 0.42** | 81.10 | **81.60 ± 0.56** |
| IFLYTEK (Acc.) | 60.25 | **60.48 ± 0.25** | 56.52 | **56.92 ± 0.17** |
| TNEWS (Acc.) | 56.24 | **56.48 ± 0.08** | 54.77 | **55.08 ± 0.16** |
| TNEWS (F1) | 55.17 | **55.39 ± 0.08** | 53.69 | **53.81 ± 0.20** |
| ChID (Acc.) | **10.66** | **10.66 ± 0.00** | **10.66** | **10.66 ± 0.00** |
| C<sup>3</sup> (Acc.) | 49.74 | **49.83 ± 0.39** | 49.42 | **50.25 ± 0.27** |

**b) Cross-Language Transfer Between Known Languages**
| GLUE Task | BERT-zh | BERT-ft-zh | mBERT | mBERT-ft-zh |
| --- | --- | --- | --- | --- |
| CoLA (MCC) | 51.13 | **53.82 ± 1.18** | 40.96 | **40.99 ± 1.08** |
| SST-2 (Acc.) | 92.26 | **92.77 ± 0.15** | 88.27 | **90.30 ± 0.66** |
| MRPC (Acc.) | 76.96 | **77.53 ± 0.28** | 82.84 | **85.20 ± 0.93** |
| MRPC (F1) | 83.38 | **84.39 ± 0.17** | 87.15 | **89.16 ± 0.78** |
| STS-B (Pears.) | 87.21 | **87.96 ± 0.12** | 87.09 | **88.30 ± 0.17** |
| STS-B (Spear.) | 86.73 | **87.67 ± 0.18** | 86.97 | **88.16 ± 0.23** |
| QQP (Acc.) | 90.65 | **90.86 ± 0.12** | 89.91 | **90.39 ± 0.13** |
| QQP (F1) | 87.53 | **87.76 ± 0.04** | 86.27 | **87.05 ± 0.09** |
| MNLI-m (Acc.) | 84.00 | **84.16 ± 0.05** | 81.52 | **81.97 ± 0.14** |
| MNLI-mm (Acc.) | 83.91 | **84.16 ± 0.10** | 81.94 | **82.78 ± 0.28** |
| QNLI (Acc.) | 91.27 | **91.42 ± 0.05** | 90.55 | **91.23 ± 0.26** |
| RTE (Acc.) | 67.15 | **67.91 ± 0.08** | 66.06 | **67.21 ± 0.67** |
| WNLI (Acc.) | 53.52 | **56.34 ± 0.00** | 54.93 | **55.25 ± 0.84** |

| CLUE Task | BERT-en | BERT-ft-en | mBERT | mBERT-ft-en |
| --- | --- | --- | --- | --- |
| AFQMC (Acc.) | 69.00 | **69.02 ± 0.03** | 69.74 | **71.25 ± 0.97** |
| CMNLI (Acc.) | 68.34 | **68.78 ± 0.13** | 78.66 | **78.87 ± 0.07** |
| CSL (F1) | 71.20 | **72.33 ± 0.66** | 81.10 | **81.51 ± 0.20** |
| IFLYTEK (Acc.) | **47.86** | 47.27 ± 0.56 | 56.52 | **56.92 ± 0.31** |
| TNEWS (Acc.) | 50.92 | **51.42 ± 0.16** | 54.77 | **54.99 ± 0.20** |
| TNEWS (F1) | 50.15 | **50.68 ± 0.07** | 53.69 | **53.76 ± 0.04** |
| ChID (Acc.) | **10.66** | **10.66 ± 0.00** | **10.66** | **10.66 ± 0.00** |
| C<sup>3</sup> (Acc.) | 42.64 | **42.99 ± 1.49** | 49.42 | **49.72 ± 0.35** |

**c) Zero-Shot Transfer to Unseen Languages**
| Task | German (de) |  | French (fr) |  | Spanish (es) |  | Japanese (ja) |  | Korean (ko) |  |
| --- | --- | --- | --- | --- | --- | --- | --- | --- | --- | --- |
|  | mBERT | mBERT-ft-en | mBERT | mBERT-ft-en | mBERT | mBERT-ft-en | mBERT | mBERT-ft-en | mBERT | mBERT-ft-en |
| XGLUE |  |  |  |  |  |  |  |  |  |  |
| PAWS-X (Acc.) | **84.00** | 83.62 ± 1.10 | **88.10** | 87.17 ± 0.81 | **87.05** | 86.50 ± 0.42 | **78.70** | 76.55 ± 2.12 | **78.75** | 76.90 ± 0.35 |
| XNLI (Acc.) | 70.60 | **70.94 ± 0.14** | 72.69 | **73.30 ± 0.80** | 74.10 | **74.62 ± 0.05** | 86.00 | **87.97 ± 0.05** | 86.00 | **87.82 ± 0.25** |
| NER (F1) | 10.67 | **12.85 ± 2.02** | 6.67 | **11.09 ± 1.09** | 7.33 | **12.11 ± 0.67** | **13.00** | 11.34 ± 2.11 | 12.00 | **12.25 ± 1.71** |
| XTREME |  |  |  |  |  |  |  |  |  |  |
| PAWS-X | **85.65** | 84.85 ± 1.34 | **88.95** | 88.00 ± 0.99 | **87.90** | 87.80 ± 0.39 | **79.55** | 78.55 ± 0.85 | **79.35** | 78.70 ± 0.64 |
| XNLI | 70.06 | **71.75 ± 0.94** | 72.97 | **73.48 ± 1.73** | 74.50 | **74.85 ± 0.35** | 86.00 | **87.61 ± 0.86** | 86.00 | **87.24 ± 0.33** |
| NER | **12.33** | 12.17 ± 1.18 | 09.33 | **13.02 ± 1.39** | 10.33 | **11.12 ± 0.30** | 08.67 | **13.00 ± 0.00** | **12.33** | 11.17 ± 1.18 |

**Table 10:** Downstream task performance before and after bilingual brain-informed fine-tuning (with language-selective). We compared vanilla (pretrained) models, English BERT (BERT-en), Chinese BERT (BERT-zh), and mBERT, with their bilingual brain-informed fine-tuned counterparts (e.g., BERT-ft-en: BERT-en fine-tuned using English brain data). For each task, the average performance and standard deviation across the six bilingual participants is reported. To assess whether brain-informed fine-tuning elicits multilingual capabilities, we evaluate downstream task performance in two settings: (a) Fine-tuning and evaluation in the same language: models are fine-tuned with brain data in one language (en or zh) and evaluated on NLP tasks in the same language (GLUE benchmark for en, CLUE benchmark for zh). This tests within-language improvements due to brain-informed fine-tuning. (b) Cross-language transfer between known languages: model is fine-tuned on brain data in one language and evaluated on tasks in the participants’s second (not used in fine-tuning) language (e.g., fine-tuned with en brain data and evaluated on CLUE (zh benchmark) tasks). This tests whether bilingual brain-informed fine-tuning elicits the participants’ shared semantic representations. Bolded values indicate performance equal to or better than the corresponding vanilla model. We observe that bilingual brain-informed fine-tuning improves performance on several NLP tasks, with mBERT showing greater benefits than BERT across all two settings. Results for fine-tuning using language-selective regions for individual participants are reported in Supplementary (see Table 5)

**a) Fine-tuning and Evaluation in the Same Language**
| GLUE Task | BERT-en | BERT-ft-en | mBERT | mBERT-ft-en |
| --- | --- | --- | --- | --- |
| CoLA (Acc.) | 53.38 | **55.25 ± 0.85** | 42.68 | 41.71 ± 0.76 |
| SST-2 (Acc.) | 92.08 | **92.62 ± 0.10** | 89.68 | **91.15 ± 0.48** |
| MRPC (Acc.) | 79.41 | **81.43 ± 0.80** | 84.80 | **85.76 ± 0.59** |
| MRPC (F1) | 86.27 | **87.47 ± 0.57** | 88.56 | **89.28 ± 0.46** |
| STS-B (Pearson) | **88.06** | 87.70 ± 0.19 | 88.06 | **88.14 ± 0.91** |
| STS-B (Spearman) | **87.65** | 87.54 ± 0.19 | 87.76 | **87.86 ± 0.78** |
| QQP (Acc.) | 90.84 | **90.88 ± 0.03** | 90.22 | **90.47 ± 0.05** |
| QQP (F1) | 87.70 | **87.79 ± 0.02** | 86.70 | **87.11 ± 0.08** |
| MNLI-m (Acc.) | **84.38** | 84.29 ± 0.23 | 82.09 | **82.10 ± 0.18** |
| MNLI-mm (Acc.) | **84.64** | 84.60 ± 0.30 | 82.38 | **82.49 ± 0.29** |
| QNLI (Acc.) | **91.45** | 91.33 ± 0.05 | 91.14 | 91.04 ± 0.11 |
| RTE (Acc.) | 67.15 | **67.47 ± 0.45** | 67.15 | **67.75 ± 1.15** |
| WNLI (Acc.) | 49.30 | **52.87 ± 0.62** | 53.52 | **56.62 ± 0.63** |

| CLUE Task | BERT-zh | BERT-ft-zh | mBERT | mBERT-ft-zh |
| --- | --- | --- | --- | --- |
| AFQMC (Acc.) | **75.25** | 73.92 ± 0.46 | 69.74 | **70.38 ± 0.50** |
| CMNLI (Acc.) | 80.50 | **80.85 ± 0.19** | 78.66 | **78.99 ± 0.26** |
| CSL (F1) | 80.18 | **81.22 ± 0.40** | 81.10 | **81.84 ± 0.27** |
| IFLYTEK (Acc.) | 60.25 | **60.41 ± 0.33** | 56.52 | **56.59 ± 0.68** |
| TNEWS (Acc.) | 56.24 | **56.49 ± 0.17** | 54.77 | **55.12 ± 0.18** |
| TNEWS (F1) | **55.17** | 55.12 ± 0.33 | 53.69 | **54.14 ± 0.19** |
| ChID (Acc.) | **10.66** | **10.66 ± 0.00** | **10.66** | **10.66 ± 0.00** |
| C<sup>3</sup> (Acc.) | 49.74 | **49.77 ± 0.33** | 49.42 | **50.15 ± 0.59** |

**b) Cross-Language Transfer Between Known Languages**
| GLUE Task | BERT-zh | BERT-ft-zh | mBERT | mBERT-ft-zh |
| --- | --- | --- | --- | --- |
| CoLA (MCC)<sup>†</sup> | 51.13 | **51.95 ± 0.56** | 40.96 | **42.50 ± 0.50** |
| SST-2 (Acc.) | 92.26 | **92.39 ± 0.13** | 88.27 | **90.89 ± 0.19** |
| MRPC (Acc.) | 76.96 | **77.74 ± 0.51** | 82.84 | **85.55 ± 0.23** |
| MRPC (F1) | 83.38 | **84.42 ± 0.89** | 87.15 | **89.35 ± 0.24** |
| STS-B (Pearson) | 87.21 | **87.63 ± 0.74** | 87.09 | **88.11 ± 0.41** |
| STS-B (Spearman) | 86.73 | **87.88 ± 0.38** | 86.97 | **88.11 ± 0.42** |
| QQP (Acc.) | 90.65 | **91.01 ± 0.31** | 89.91 | **90.44 ± 0.05** |
| QQP (F1) | 87.53 | **87.90 ± 0.21** | 86.27 | **87.07 ± 0.06** |
| MNLI-m (Acc.) | 84.00 | **84.19 ± 0.42** | 81.52 | **82.09 ± 0.17** |
| MNLI-mm (Acc.) | 83.91 | **84.17 ± 0.47** | 81.94 | **82.47 ± 0.34** |
| QNLI (Acc.) | 91.27 | **91.33 ± 0.10** | 90.55 | **91.12 ± 0.19** |
| RTE (Acc.) | **67.15** | 66.81 ± 0.67 | 66.06 | **66.63 ± 1.28** |
| WNLI (Acc.) | 53.52 | **54.38 ± 1.88** | 54.93 | **55.49 ± 1.26** |

| CLUE Task | BERT-en | BERT-ft-en | mBERT | mBERT-ft-en |
| --- | --- | --- | --- | --- |
| AFQMC (Acc.) | 69.00 | **69.31 ± 0.47** | 69.74 | **70.78 ± 0.67** |
| CMNLI (Acc.) | 68.34 | **68.34 ± 0.63** | 78.66 | **78.88 ± 0.09** |
| CSL (F1) | 71.20 | **71.46 ± 0.43** | 81.10 | **81.25 ± 0.29** |
| IFLYTEK (Acc.) | 47.86 | **48.05 ± 1.44** | 56.52 | **57.02 ± 0.29** |
| TNEWS (Acc.) | **50.92** | 50.69 ± 0.18 | 54.77 | **55.23 ± 0.24** |
| TNEWS (F1) | 50.15 | **50.48 ± 0.18** | 53.69 | **54.11 ± 0.11** |
| ChID (Acc.) | **10.66** | **10.66 ± 0.00** | **10.66** | **10.66 ± 0.00** |
| C<sup>3</sup> (Acc.) | 42.64 | **55.18 ± 0.61** | 49.42 | **50.24 ± 0.33** |

Detailed results for individual participants, including fine-tuning with whole-brain, language-selective, and semantically-selective regions, are provided in the Supplementary (Tables 4–6). Results and analyses for other multilingual language models (XLM-R, XGLM, and LLaMA) are also reported in the Supplementary.

### 4.3 Monolingual Brain data as Fine-Tuning Targets

Table 2 compares downstream task performance after brain-informed fine-tuning (with semantically-selective brain regions) with English brain data from either bilingual or monolingual participants. Monolingual brain-informed fine-tuning improves downstream performance relative to the baseline across several tasks. However, bilingual fine-tuning yields even greater gains. On the GLUE bench-mark (within-language evaluation), bilingual brain-informed fine-tuning outperforms monolingual tuning on 7/9 tasks, especially in all inference-related tasks (MNLI, QNLI, and WNLI). On the CLUE benchmark (cross-language evaluation), bilingual brain-informed fine-tuning leads to higher performance on 5/7 tasks. These results suggest that while monolingual brain-informed fine-tuning also improves downstream tasks within-language, fine-tuning with bilingual brain data enhances cross-linguistic generalization. Results for fine-tuning using whole-brain brain data are reported in Appendix Table 11.

## 5 Discussion and Conclusion

In this work, we perform brain-informed fine-tuning of monolingual and multilingual language models using brain data from bilingual participants reading the same naturalistic stories in English and Chinese. We show that bilingual brain-informed fine-tuned language models outperform their vanilla (pretrained) counterparts in both brain encoding performance and most downstream NLP tasks. Our analysis of brain-informed fine-tuning reveals several key conclusions.

First, representations from brain-informed fine-tuned language models are shared across bilingual individuals. Both monolingual and multilingual language models show improved encoding performance after brain-informed fine-tuning, in both English and Chinese (Fig.2). These models outperform baseline controls (Appendix Fig.4) and generalize across participants (Appendix Fig.6). This highlights the potential of bilingual brain data as a rich supervisory signal for grounding language model representations with biologically meaningful representations. Second, brain-informed fine-tuning enables cross-language transfer in language models. Downstream performance improves across within-language, cross-language, and unseen language settings (Table 1). This holds for fine-tuning with whole-brain or language-selective or semantically-selective regions (Appendix Tables 9 and 8). This effect is likely driven by shared semantic representations in bilingual individuals (Chen et al., 2024b). This highlights the potential of leveraging bilingual brain representations for developing language-agnostic language models. Lastly, the observed improvements are driven specifically by brain-informed fine-tuning with bilingual brain data, not brain data in general. Fine-tuning with monolingual brain data does not yield similar cross-linguistic transfer (Table 2).

Our results contribute to a growing body of work exploring the representational alignment between brain and artificial language models. We demonstrate that bilingual brain-informed fine-tuning improves multilingual understanding in language models. To the best of our knowledge, this is the first work to do so. This study offers a bridge between neuroscience research on cross-linguistic semantic representation and NLP efforts to interpret and improve multilingual language models.

## 6 Limitations and Future Work

A key limitation of this study is the relatively small number of data samples per participant, due to limited bilingual brain recordings with naturalistic stimuli. This constrains the observed performance gains. Additionally, our analysis is restricted to only two languages (English and Chinese). Expanding to larger and more diverse multilingual brain datasets would allow a broader evaluation of cross-linguistic transfer. Future work could also examine what linguistic properties are captured by brain-informed fine-tuning (such as syntax, morphology, or discourse-level structure). This can potentially be tested by targeting functionally selective brain regions and analyzing fine-tuned model representations across layers.

## Acknowledgment

We thank Lily Gong for collecting and preprocessing the fMRI data, and Mathis Lamarre for valuable discussions. This work was funded by grants from the German Federal Ministry of Education and Research (BMBF; Grant no. 01GQ1906) and the European Research Council (ERC; Grant no. 101042567).

## NeurIPS Paper Checklist

### 1. Claims

Question: Do the main claims made in the abstract and introduction accurately reflect the paper’s contributions and scope?

Answer: [Yes]

Justification: We have ensured that the main claims made in the abstract and introduction are directly correlating to the research findings and the methods we have employed.

Guidelines:

- The answer NA means that the abstract and introduction do not include the claims made in the paper.
- The abstract and/or introduction should clearly state the claims made, including the contributions made in the paper and important assumptions and limitations. A No or NA answer to this question will not be perceived well by the reviewers.
- The claims made should match theoretical and experimental results, and reflect how much the results can be expected to generalize to other settings.
- It is fine to include aspirational goals as motivation as long as it is clear that these goals are not attained by the paper.

### 2. Limitations

Question: Does the paper discuss the limitations of the work performed by the authors? Answer: [Yes]

Justification: The paper discusses the main limitations of the work performed by the authors in the discussion section.

Guidelines:

- The answer NA means that the paper has no limitation while the answer No means that the paper has limitations, but those are not discussed in the paper.
- The authors are encouraged to create a separate “Limitations” section in their paper.
- The paper should point out any strong assumptions and how robust the results are to violations of these assumptions (e.g., independence assumptions, noiseless settings, model well-specification, asymptotic approximations only holding locally). The authors should reflect on how these assumptions might be violated in practice and what the implications would be.
- The authors should reflect on the scope of the claims made, e.g., if the approach was only tested on a few datasets or with a few runs. In general, empirical results often depend on implicit assumptions, which should be articulated.
- The authors should reflect on the factors that influence the performance of the approach. For example, a facial recognition algorithm may perform poorly when image resolution is low or images are taken in low lighting. Or a speech-to-text system might not be used reliably to provide closed captions for online lectures because it fails to handle technical jargon.
- The authors should discuss the computational efficiency of the proposed algorithms and how they scale with dataset size.
- If applicable, the authors should discuss possible limitations of their approach to address problems of privacy and fairness.
- While the authors might fear that complete honesty about limitations might be used by reviewers as grounds for rejection, a worse outcome might be that reviewers discover limitations that aren’t acknowledged in the paper. The authors should use their best judgment and recognize that individual actions in favor of transparency play an important role in developing norms that preserve the integrity of the community. Reviewers will be specifically instructed to not penalize honesty concerning limitations.

### 3. Theory assumptions and proofs

Question: For each theoretical result, does the paper provide the full set of assumptions and a complete (and correct) proof?

Answer: [NA]

Justification: Our paper does not require any explicit theorems and proofs. Guidelines:

- The answer NA means that the paper does not include theoretical results.
- All the theorems, formulas, and proofs in the paper should be numbered and cross-referenced.
- All assumptions should be clearly stated or referenced in the statement of any theorems.
- The proofs can either appear in the main paper or the supplemental material, but if they appear in the supplemental material, the authors are encouraged to provide a short proof sketch to provide intuition.
- Inversely, any informal proof provided in the core of the paper should be complemented by formal proofs provided in appendix or supplemental material.
- Theorems and Lemmas that the proof relies upon should be properly referenced.

### 4. Experimental result reproducibility

Question: Does the paper fully disclose all the information needed to reproduce the main experimental results of the paper to the extent that it affects the main claims and/or conclusions of the paper (regardless of whether the code and data are provided or not)?

Answer: [Yes]

Justification: The paper has delineated all the information related to the experimental setup in the experimental setup section and model parameters details in Appendix D.

Guidelines:

- The answer NA means that the paper does not include experiments.
- If the paper includes experiments, a No answer to this question will not be perceived well by the reviewers: Making the paper reproducible is important, regardless of whether the code and data are provided or not.
- If the contribution is a dataset and/or model, the authors should describe the steps taken to make their results reproducible or verifiable.
- Depending on the contribution, reproducibility can be accomplished in various ways. For example, if the contribution is a novel architecture, describing the architecture fully might suffice, or if the contribution is a specific model and empirical evaluation, it may be necessary to either make it possible for others to replicate the model with the same dataset, or provide access to the model. In general, releasing code and data is often one good way to accomplish this, but reproducibility can also be provided via detailed instructions for how to replicate the results, access to a hosted model (e.g., in the case of a large language model), releasing of a model checkpoint, or other means that are appropriate to the research performed.
- While NeurIPS does not require releasing code, the conference does require all submissions to provide some reasonable avenue for reproducibility, which may depend on the nature of the contribution. For example
  a. If the contribution is primarily a new algorithm, the paper should make it clear how to reproduce that algorithm.
  b. If the contribution is primarily a new model architecture, the paper should describe the architecture clearly and fully.
  c. If the contribution is a new model (e.g., a large language model), then there should either be a way to access this model for reproducing the results or a way to reproduce the model (e.g., with an open-source dataset or instructions for how to construct the dataset).
  d. We recognize that reproducibility may be tricky in some cases, in which case authors are welcome to describe the particular way they provide for reproducibility. In the case of closed-source models, it may be that access to the model is limited in some way (e.g., to registered users), but it should be possible for other researchers to have some path to reproducing or verifying the results.

### 5. Open access to data and code

Question: Does the paper provide open access to the data and code, with sufficient instructions to faithfully reproduce the main experimental results, as described in supplemental material?

Answer: [Yes]

Justification: We provide a GitHub repository in the abstract. Guidelines:

- The answer NA means that paper does not include experiments requiring code.
- Please see the NeurIPS code and data submission guidelines (https://nips.cc/public/guides/CodeSubmissionPolicy) for more details.
- While we encourage the release of code and data, we understand that this might not be possible, so “No” is an acceptable answer. Papers cannot be rejected simply for not including code, unless this is central to the contribution (e.g., for a new open-source benchmark).
- The instructions should contain the exact command and environment needed to run to reproduce the results. See the NeurIPS code and data submission guidelines (https://nips.cc/public/guides/CodeSubmissionPolicy) for more details.
- The authors should provide instructions on data access and preparation, including how to access the raw data, preprocessed data, intermediate data, and generated data, etc.
- The authors should provide scripts to reproduce all experimental results for the new proposed method and baselines. If only a subset of experiments are reproducible, they should state which ones are omitted from the script and why.
- At submission time, to preserve anonymity, the authors should release anonymized versions (if applicable).
- Providing as much information as possible in supplemental material (appended to the paper) is recommended, but including URLs to data and code is permitted.

### 6. Experimental setting/details

Question: Does the paper specify all the training and test details (e.g., data splits, hyperparameters, how they were chosen, type of optimizer, etc.) necessary to understand the results?

Answer: [Yes]

Justification: We provide all the training and test details in the experimental setup (See Section 3.3, 3.4 and 3.5).

Guidelines:

- The answer NA means that the paper does not include experiments.
- The experimental setting should be presented in the core of the paper to a level of detail that is necessary to appreciate the results and make sense of them.
- The full details can be provided either with the code, in appendix, or as supplemental material.

### 7. Experiment statistical significance

Question: Does the paper report error bars suitably and correctly defined or other appropriate information about the statistical significance of the experiments?

Answer: [Yes]

Justification: We conducted our experiments multiple times across 6 participants and took the average results for baseline settings.

Guidelines:

- The answer NA means that the paper does not include experiments.
- The authors should answer “Yes” if the results are accompanied by error bars, confidence intervals, or statistical significance tests, at least for the experiments that support the main claims of the paper.
- The factors of variability that the error bars are capturing should be clearly stated (for example, train/test split, initialization, random drawing of some parameter, or overall run with given experimental conditions).
- The method for calculating the error bars should be explained (closed form formula, call to a library function, bootstrap, etc.)
- The assumptions made should be given (e.g., Normally distributed errors).
- It should be clear whether the error bar is the standard deviation or the standard error of the mean.
- It is OK to report 1-sigma error bars, but one should state it. The authors should preferably report a 2-sigma error bar than state that they have a 96% CI, if the hypothesis of Normality of errors is not verified.
- For asymmetric distributions, the authors should be careful not to show in tables or figures symmetric error bars that would yield results that are out of range (e.g. negative error rates).
- If error bars are reported in tables or plots, The authors should explain in the text how they were calculated and reference the corresponding figures or tables in the text.

### 8. Experiments compute resources

Question: For each experiment, does the paper provide sufficient information on the computer resources (type of compute workers, memory, time of execution) needed to reproduce the experiments?

Answer: [Yes]

Justification: We have included the specifications of the hardware and software environments to ensure the reproducibility of our results (See Appendix D).

Guidelines:

- The answer NA means that the paper does not include experiments.
- The paper should indicate the type of compute workers CPU or GPU, internal cluster, or cloud provider, including relevant memory and storage.
- The paper should provide the amount of compute required for each of the individual experimental runs as well as estimate the total compute.
- The paper should disclose whether the full research project required more compute than the experiments reported in the paper (e.g., preliminary or failed experiments that didn’t make it into the paper).

### 9. Code of ethics

Question: Does the research conducted in the paper conform, in every respect, with the NeurIPS Code of Ethics https://neurips.cc/public/EthicsGuidelines?

Answer: [Yes]

Justification: The research conducted in this paper fully conforms with the NeurIPS Code of Ethics in every respect.

Guidelines:

- The answer NA means that the authors have not reviewed the NeurIPS Code of Ethics.
- If the authors answer No, they should explain the special circumstances that require a deviation from the Code of Ethics.
- The authors should make sure to preserve anonymity (e.g., if there is a special consideration due to laws or regulations in their jurisdiction).

### 10. Broader impacts

Question: Does the paper discuss both potential positive societal impacts and negative societal impacts of the work performed?

Answer: [Yes]

Justification: This paper could benefit society by advancing the development of NLP systems to be more human-like.

Guidelines:

- The answer NA means that there is no societal impact of the work performed.
- If the authors answer NA or No, they should explain why their work has no societal impact or why the paper does not address societal impact.
- Examples of negative societal impacts include potential malicious or unintended uses (e.g., disinformation, generating fake profiles, surveillance), fairness considerations (e.g., deployment of technologies that could make decisions that unfairly impact specific groups), privacy considerations, and security considerations.
- The conference expects that many papers will be foundational research and not tied to particular applications, let alone deployments. However, if there is a direct path to any negative applications, the authors should point it out. For example, it is legitimate to point out that an improvement in the quality of generative models could be used to generate deepfakes for disinformation. On the other hand, it is not needed to point out that a generic algorithm for optimizing neural networks could enable people to train models that generate Deepfakes faster.
- The authors should consider possible harms that could arise when the technology is being used as intended and functioning correctly, harms that could arise when the technology is being used as intended but gives incorrect results, and harms following from (intentional or unintentional) misuse of the technology.
- If there are negative societal impacts, the authors could also discuss possible mitigation strategies (e.g., gated release of models, providing defenses in addition to attacks, mechanisms for monitoring misuse, mechanisms to monitor how a system learns from feedback over time, improving the efficiency and accessibility of ML).

### 11. Safeguards

Question: Does the paper describe safeguards that have been put in place for responsible release of data or models that have a high risk for misuse (e.g., pretrained language models, image generators, or scraped datasets)?

Answer: [NA]

Justification: Our research does not pose any risks for misuse. Guidelines:

- The answer NA means that the paper poses no such risks.
- Released models that have a high risk for misuse or dual-use should be released with necessary safeguards to allow for controlled use of the model, for example by requiring that users adhere to usage guidelines or restrictions to access the model or implementing safety filters.
- Datasets that have been scraped from the Internet could pose safety risks. The authors should describe how they avoided releasing unsafe images.
- We recognize that providing effective safeguards is challenging, and many papers do not require this, but we encourage authors to take this into account and make a best faith effort.

### 12. Licenses for existing assets

Question: Are the creators or original owners of assets (e.g., code, data, models), used in the paper, properly credited and are the license and terms of use explicitly mentioned and properly respected?

Answer: [Yes]

Justification: We have explicitly cited the NLP benchmark datasets, code, and models used. Guidelines:

- The answer NA means that the paper does not use existing assets.
- The authors should cite the original paper that produced the code package or dataset.
- The authors should state which version of the asset is used and, if possible, include a URL.
- The name of the license (e.g., CC-BY 4.0) should be included for each asset.
- For scraped data from a particular source (e.g., website), the copyright and terms of service of that source should be provided.
- If assets are released, the license, copyright information, and terms of use in the package should be provided. For popular datasets, paperswithcode.com/datasets has curated licenses for some datasets. Their licensing guide can help determine the license of a dataset.
- For existing datasets that are re-packaged, both the original license and the license of the derived asset (if it has changed) should be provided.
- If this information is not available online, the authors are encouraged to reach out to the asset’s creators.

### 13. New assets

Question: Are new assets introduced in the paper well documented and is the documentation provided alongside the assets?

Answer: [Yes]

Justification: We release the code, model weights, and complete documentation in the GitHub repository linked in the abstract.

Guidelines:

- The answer NA means that the paper does not release new assets.
- Researchers should communicate the details of the dataset/code/model as part of their submissions via structured templates. This includes details about training, license, limitations, etc.
- The paper should discuss whether and how consent was obtained from people whose asset is used.
- At submission time, remember to anonymize your assets (if applicable). You can either create an anonymized URL or include an anonymized zip file.

### 14. Crowdsourcing and research with human subjects

Question: For crowdsourcing experiments and research with human subjects, does the paper include the full text of instructions given to participants and screenshots, if applicable, as well as details about compensation (if any)?

Answer: [Yes]

Justification: fMRI data was recorded from human subjects. All ethical regulations relevant to human research participants were followed, and all procedures were approved by the Committee for the Protection of Human Subjects.

Guidelines:

- The answer NA means that the paper does not involve crowdsourcing nor research with human subjects.
- Including this information in the supplemental material is fine, but if the main contribution of the paper involves human subjects, then as much detail as possible should be included in the main paper.
- According to the NeurIPS Code of Ethics, workers involved in data collection, curation, or other labor should be paid at least the minimum wage in the country of the data collector.

### 15. Institutional review board (IRB) approvals or equivalent for research with human subjects

Question: Does the paper describe potential risks incurred by study participants, whether such risks were disclosed to the subjects, and whether Institutional Review Board (IRB) approvals (or an equivalent approval/review based on the requirements of your country or institution) were obtained?

Answer: [Yes]

Justification: All subjects signed a consent form where risks were disclosed, and IRB approval was obtained for this experiment. Further details can be found in Appendix B.2.

Guidelines:

- The answer NA means that the paper does not involve crowdsourcing nor research with human subjects.
- Depending on the country in which research is conducted, IRB approval (or equivalent) may be required for any human subjects research. If you obtained IRB approval, you should clearly state this in the paper.
- We recognize that the procedures for this may vary significantly between institutions and locations, and we expect authors to adhere to the NeurIPS Code of Ethics and the guidelines for their institution.
- For initial submissions, do not include any information that would break anonymity (if applicable), such as the institution conducting the review.

### 16. Declaration of LLM usage

Question: Does the paper describe the usage of LLMs if it is an important, original, or non-standard component of the core methods in this research? Note that if the LLM is used only for writing, editing, or formatting purposes and does not impact the core methodology, scientific rigorousness, or originality of the research, declaration is not required.

Answer: [Yes]

Justification: We have used LLM only for grammar correction. Guidelines:

- The answer NA means that the core method development in this research does not involve LLMs as any importantss, original, or non-standard components.
- Please refer to our LLM policy (https://neurips.cc/Conferences/2025/LLM) for what should or should not be described.

## Overview of Appendices

- Section A: Detailed Related Work
- Section B: Naturalistic Brain Imaging Dataset and Preprocessing
- Section C: Brain-Informed Fine-tuning with Subsets of Cortex
- Section D: Implementation details for reproducibility
- Section E: Details of Downstream NLP Tasks
- Section F: Additional results for Voxelwise encoding performance
- Section G Models fine-tuned with different training objectives
- Section H Bilingual vs. Monolingual with English whole-brain data

## A. Detailed Related Work

### Fine-tuning of language models with naturalistic brain data

Our work is most closely related to that of Schwartz et al. (2019); Moussa et al. (2025); Vattikonda et al. (2025), who proposed the brain-tuning approach to fine-tune pretrained Transformer-based language models using brain data. This approach integrates brain-relevant information into the language models and examines whether and how brain-informed language models effect brain encoding performance and downstream task performance. Specifically, Schwartz et al. (2019) introduced brain-tuning using brain data recorded during naturalistic reading tasks in monolingual English speakers, observing improved brain encoding performance after fine-tuning along with enhanced performance on downstream NLP tasks. More recently, Moussa et al. (2025); Vattikonda et al. (2025) applied a similar approach to explore whether brain-tuning of speech-based language models could enhance brain-relevant semantic representations, thus improving encoding performance and speech-task performance. Our study complements these previous studies by investigating bilingual brain-informed fine-tuning and exploring how monolingual and multilingual language models change when trained with bilingual brain data.

### Multilingual language models and brain alignment

Our work also relates to a growing body of literature investigating alignment between human brain activity and language models. Several studies have successfully used text-based language models to predict brain activity evoked by both written and spoken stimuli, achieving impressive levels of alignment between language models and brain activity (Wehbe et al., 2014a; Jain & Huth, 2018; Toneva & Wehbe, 2019; Deniz et al., 2019; Abdou et al., 2021; Toneva et al., 2022; Antonello et al., 2021; Oota et al., 2022; Aw & Toneva, 2023; Oota et al., 2024b; Lamarre et al., 2022; Chen et al., 2024a). More recently, research has focused on multilingual Transformer-based language models using brain data from reading and listening tasks across multiple languages to evaluate their brain encoding performance (de Varda et al., 2025). However, previous multilingual studies have generally remained monolingual in their experimental design, with participants exposed exclusively to stimuli in one language. A notable exception is the recent work by Chen et al. (2024a), which examined bilingual language processing in participants who read identical stories in both English and Chinese, and find that bilingual individuals have shared semantic representations across languages. Our approach complements this research by further exploring bilingual brain alignment through brain-informed fine-tuning of language models, particularly analyzing how different brain regions respond to bilingual stimuli and fine-tuning strategies, evaluating their impact on downstream NLP tasks in both English and Chinese.

## B Naturalistic Brain Imaging Dataset and Preprocessing

### B.1 MRI data collection of the bilingual dataset

MRI data were collected on a 3T Siemens TIM Trio scanner using a 32-channel volume coil. Functional scans used a gradient-echo EPI sequence (TR = 2.0045 s, TE = 35 ms, flip angle = 74°, voxel size = 2.24 × 2.24 × 4.1 mm, 30 interleaved axial slices). Motion correction and alignment were performed using the FMRIB Linear Image Registration Tool (FLIRT) from FSL. Please refer to Chen et al. (2024b) for more details.

### B.2 Participants

Functional data from six healthy bilingual (in English and Chinese) participants from (Chen et al., 2024b) with normal hearing and vision was used. All participants were right-handed or ambidextrous. All potential risks and/or discomforts from MRI were disclosed to participants: 1) loud beeping and hammering sounds while the scanner is collecting measurements; 2) claustrophobia while inside the scanner; 3) nerve stimulation; 4) risk of injury from metal objects affected by the scanner (highly unlikely); 5) compromised confidentiality (highly unlikely). All participants signed a consent form where these risks were disclosed. All ethical regulations relevant to human research participants were followed, and all procedures were approved by the Committee for the Protection of Human Participants. For anonymity, detailed information about the ethics committee has been omitted and will be included in the camera-ready version. Voxelwise encoding models and brain-informed fine-tuning are done separately for each participant, with results reported per participant. No sample size calculations were performed, as each participant serves as a full replication of the results.

### B.3 Monolingual participants from LeBel et al. (2023) and Deniz et al. (2019)

Functional data were collected from three healthy, monolingual English participants with normal hearing: two participants (UTS07: male, age 25; UTS08: male, age 24) from LeBel et al. (2023) and one participant (male) from (Deniz et al., 2019). Participants listened to several narrative stories from *The Moth* podcast. For the monolingual analysis, we used a subset of the data from LeBel et al. (2023) and Deniz et al. (2019), specifically the same stories that are included in the bilingual dataset. For more details, please refer to the original publications (Deniz et al., 2019; LeBel et al., 2023).

## C Brain-Informed Fine-tuning with Subsets of Cortex

### C.1 Fine-tuning with Language-Selective Regions

Voxels were extracted from regions of interest (ROIs) known to support language comprehension (Fedorenko et al., 2010). To apply these ROIs to individual participants, we first mapped each participant’s cortical surface to the fsaverage template using surface-based registration. The group-level language mask was then projected onto each participant’s brain to extract participant-specific language ROIs. The ROI mask used is shown in Appendix Fig. 3a.

### C.2 Fine-tuning with Semantically-Selective Regions

To identify voxels selective for semantics, we applied a two-stage regression procedure to isolate semantic signals while controlling for low-level sensory confounds. Semantic features were extracted using fastText (Bojanowski et al., 2017) embeddings of the narrative text. This procedure was performed separately for each participant and language. First, we trained VEMs for response to seven low-level sensory feature spaces: word count, letter/character count, single phonemes, diphones, triphones, visual motion energy, and pixel-based orthographic similarity. These models were trained on the training data and used to predict responses for both train and test sets. Predicted low-level responses were subtracted from the actual BOLD signals to obtain residuals. These residual responses were then z-scored per voxel to have unit variance. Next, we trained VEMs using the semantic features on the residual signals. Voxels that showed significant (one-sided permutation test, p < 0.05, FDR-corrected) prediction accuracy based on semantic features were labeled as semantically-selective. The semantically-selective voxel mask for participant 1 is shown in Appendix Fig. 3b.

### C.3 Baseline Models for Fine-tuning

#### TR-shuffled fMRI as fine-tuning target

We tested how fine-tuning language models with temporally misaligned data affects model representation. This TR-shuffled approach allows us to verify that the improvements from brain-informed fine-tuning are driven by meaningful stimulus-response alignment, rather than random brain data. Here, we randomly shuffled the brain responses, permuted in blocks of 10 contiguous TRs. This breaks the correspondence between stimulus and brain response.

#### Multilingual model features as fine-tuning target

We tested how fine-tuning monolingual language models with multilingual model representations affects model representations. The goal of this test is to compare the results between pure-language model fine-tuning with brain-informed language model fine-tuning. Here, we replaced the fMRI data associated with input stimuli with representations obtained from a multilingual language model. We then fine-tuned monolingual language models (BERT-en and BERT-zh) using these multilingual model representations (from mBERT) as targets.

#### Monolingual brain data as fine-tuning target

We tested how tuning language models with brain data from individuals who know only one language differs from tuning language models with bilingual brain data. This test is used to clarify the impact of bilingualism on the resulting model representations.

## D Implementation details for reproducibility

All brain-informed fine-tuning experiments were conducted on a machine equipped with an NVIDIA TITAN RTX GPU (24 GB RAM), and NVIDIA RTX A6000 GPU (40GB RAM). The downstream NLP tasks were evaluated on machines with the same GPU configuration. Voxelwise encoding models were trained on the TITAN RTX GPU using banded ridge regression, with the following hyperparameters: MSE loss, L2 regularization (*λ*) values logarithmically spaced from 10^*−*10^ to 10^10^, target batch size of 1000, and 20 *λ* values per batch.

For downstream NLP task fine-tuning, we used a batch size of 64, learning rate of 2e-5, weight decay of 0.01, per-device evaluation batch size of 128, and trained for 3 epochs.

## E Details of Downstream NLP Tasks

Detailed description of the downstream NLP tasks in GLUE and CLUE benchmarks is provided in Table 3. Detailed description of the downstream NLP tasks in XGLUE and XTREME benchmarks is provided in Table 4. The metric ranges and the scaling for each task are provided in Tables 5, 6, and 7.

## F Additional results for Voxelwise Encoding Performance

Please note that the bilingual dataset is from a reading experiment, where variations in visual brain areas is correlated with stimulus presentation, especially in naturalistic settings. Moreover, semantic features from language models are known to spuriously predict brain activity in visual areas even after regressing out low-level visual information (Deniz et al., 2019; Oota et al., 2024b). In our analyses, we used embeddings extracted from the language model layer for fitting the VEMs, without including any low-level sensory features. Consequently, well-predicted voxels in the reported figures do not necessarily correspond to language-selective brain regions. This distinction is important to avoid overinterpreting encoding performance in non-linguistic areas such as the early visual cortex. Importantly, our analyses using semantically selective voxels (which explicitly exclude visual regions) show consistent performance gains from brain-informed fine-tuning, indicating that the observed improvements are not driven by visual areas but reflect genuine alignment with higher-level semantic processing.

### F.1 Baseline Models

Appendix Fig. 4 shows results from baseline experiments where brain-informed fine-tuning was performed using (a) TR-shuffled fMRI responses and (b) mBERT representations as fine-tuning targets. For BERT-en across participants (considering well-predicted voxels with *r >* 0.1), the average difference in encoding performance was 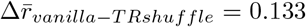 and 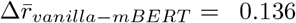. In both cases, we observe that VEM predictions are limited to some sensory regions and scattered across the cortex, with no systematic predictions in higher-level semantic areas. This confirms that the observed improvements in our main experiments are not driven by spurious correlations but rather due to biologically plausible representations in bilingual brains.

### F.2 Results on Participants 2-6

Fig. 5 shows brain encoding performance before and after brain-informed fine-tuning for all participants. This figure shows voxelwise encoding model performance for the best-performing fine-tuned variants across the cortex for all participants, using (a) BERT-en, (b) BERT-zh, and (c–d) mBERT models. This replicates the format of Fig. 2, and reports it for all participants. For details on voxel color coding, refer to the caption of Fig. 2. Across participants, model types, and languages, brain-informed fine-tuning consistently improves encoding performance relative to it’s vanilla counterpart.

### F.3 Cross-participant Transfer

To examine whether brain-informed fine-tuning generalizes encoding performance across participants, we evaluated brain-informed fine-tuned models on one participant’s brain data and test it on the brain recordings of other participants. Fig. 6 presents the encoding performance comparisons between vanilla and brain-tuned models in this cross-participant setting. Across participants (considering well-predicted voxels with *r >* 0.1), the average difference in encoding performance was 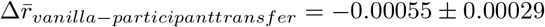.

We observe that there is also no difference in overall encoding performance between the vanilla and brain-tuned models when tested on unseen participants. This pattern holds consistently across all participants, for both English and Chinese brain data, and for both monolingual and multilingual BERT models. These findings suggest that the representational changes introduced by brain-specific fine-tuning are not participant-specific but reflect some shared representations across bilingual individuals.

## G Additional results on Downstream NLP Tasks Performance

### G.1 Vanilla vs. Fine-tuned with Whole-brain/Language-selective regions/Semantically-selective voxels

Table 8 shows downstream task performance before and after bilingual brain-informed fine-tuning (with whole-brain, language-selective, and semantically-selective voxels). We compared vanilla (pretrained) models and mBERT, with their bilingual brain-informed fine-tuned counterparts (e.g., mBERT-ft-en and mBERT-ft-zh). To assess whether brain-informed fine-tuning elicit multilingual capabilities, we evaluate downstream task performance in two settings: (a) Fine-tuning and Evaluation in the Same Language: models are fine-tuned with brain data in one language (English (en) or Chinese (zh)) and evaluated on NLP tasks in the same language (GLUE benchmark for en, CLUE benchmark for zh). This tests within-language improvements due to brain-informed fine-tuning. (b) Cross-Language Transfer Between Known Languages: model is fine-tuned on brain data in one language and evaluated on tasks in the participants’s second (not used in fine-tuning) language (e.g., fine-tuned with en brain data and evaluated on CLUE (zh benchmark) task. This tests whether bilingual brain-informed fine-tuning elicits the participants’ shared semantic representations. Bolded values indicate equal or the best performance. We observe that bilingual brain-informed fine-tuning improves performance on several NLP tasks, with either language- or semantically-selective region variants showing more improvement than their vanilla counterpart.

## H Bilingual vs. Monolingual with English whole-brain data

**Table 11:** Downstream task performance before and after bilingual or monolingual brain-informed fine-tuning. We perform brain-informed fine-tuning of mBERT with English whole-brain brain data (mBERT-ft-en) from either a bilingual (participant 1) or a monolingual participant. We evaluate downstream task performance in two settings: (a) Fine-tuning and evaluation in the same language: the model is evaluated in English with GLUE tasks. (b) Cross-language transfer between known languages: the model is evaluated in Chinese with CLUE tasks.

**a) Fine-tuning and Evaluation in the Same Language**
| Task | mBERT | mBERT-ft-en |  |
| --- | --- | --- | --- |
|  |  | Bilingual | Monolingual |
| CoLA (MCC) | **42.68** | 40.05 | 35.74 |
| SST-2 (Acc.) | 89.68 | 90.14 | **90.48** |
| MRPC (Acc.) | 84.80 | **84.80** | 78.85 |
| MRPC (F1) | 88.56 | **89.01** | 85.78 |
| STS-B (Pearson) | 88.06 | **88.42** | **88.42** |
| STS-B (Spearman) | 87.76 | 88.22 | **88.34** |
| QQP (Acc.) | 90.22 | 90.47 | **90.54** |
| QQP (F1) | 86.70 | 87.07 | **87.19** |
| MNLI-m (Acc.) | 82.09 | **82.66** | 82.41 |
| MNLI-mm (Acc.) | 82.38 | **82.97** | 82.78 |
| QNLI (Acc.) | 91.14 | **91.18** | 91.12 |
| RTE (Acc.) | 67.15 | **65.70** | 65.34 |
| WNLI (Acc.) | 53.52 | **56.34** | **56.34** |

**b) Cross-Language Transfer Between Known Languages**
| CLUE Task | mBERT | mBERT-ft-en |  |
| --- | --- | --- | --- |
|  |  | Bilingual | Monolingual |
| AFQMC (Acc.) | 69.74 | 70.62 | **70.64** |
| CMNLI (Acc.) | 78.66 | **78.83** | 78.48 |
| CSL (F1) | 81.10 | **81.30** | 81.18 |
| IFLYTEK (Acc.) | 46.79 | 56.52 | **57.06** |
| TNEWS (Acc.) | 54.77 | **54.81** | 54.80 |
| TNEWS (F1) | 53.69 | **53.84** | 53.50 |
| ChID (Acc.) | 10.66 | **10.66** | **10.66** |
| C<sup>3</sup> (Acc.) | 49.42 | **49.48** | 49.46 |

## Footnotes

2 https://github.com/denizenslab/brain-informed-fine-tuning

3 *”TR” stands for repetition time, which is the acquisition time for each fMRI volume; in our case, the TR was 2*.*0045 seconds*.

