## Supplementary material for "Brain-Informed Fine-Tuning for Improved Multilingual Understanding in Language Models": NeurIPS_2025_Bling_Tuning_Supplemenatry.pdf

### Overview of Supplementary

- Section 1: Results with Alternative Loss Functions
- Section 2: Comparison of Multilingual Language Models
- Section 3: Results with LLaMA-3.2-1B
- Section 4: Performance on Downstream NLP Tasks for Individual Participants.
- Section 5: Comparison with a Previously Proposed Fine-Tuning Pipeline
- Section 6: Scaling Effects on Encoding Performance in Brain-Informed Fine-Tuning

#### 1 Results with Alternative Loss Functions

We would like to note that contrastive loss yielded better brain encoding performance (mean r = 0.163, SE = 0.0097), slightly outperforming Ridge (mean r = 0.1622) and MSE (mean r = 0.1612). Given that contrastive objectives are commonly used in prior brain decoding studies and offer strong alignment with brain representations (Défossez et al., 2023; Lévy et al., 2025), we retained it as our main objective to ensure comparability with existing literature.

Table 1: **Downstream task performance before and after bilingual brain-informed fine-tuning (with whole-brain) with different loss functions.** We compared vanilla (pretrained) mBERT model with their bilingual brain-informed fine-tuned counterparts (e.g., mBERT-ft-en: mBERT fine-tuned using English brain data) with different loss functions used during training. We evaluate downstream task performance in two settings: (a) Fine-tuning and evaluation in the same language: models are fine-tuned with brain data in one language (en or zh) and evaluated on NLP tasks in the same language (GLUE benchmark for en, CLUE benchmark for zh). This tests within-language improvements due to brain-informed fine-tuning. (b) Cross-language transfer between known languages: model is fine-tuned on brain data in one language and evaluated on tasks in the participants' second (not used in fine-tuning) language (e.g., fine-tuned with en brain data and evaluated on CLUE (zh benchmark) tasks). This tests whether bilingual brain-informed fine-tuning elicits the participants' shared semantic representations. Bolded values indicate performance equal to or better than the corresponding vanilla model.

**a) Fine-tuning and Evaluation in the Same Language**

| | | mBERT-ft-en | | | | |
|---|---|---|---|---|---|---|
| **GLUE Task** | **mBERT** | **Contrastive** | **MSE** | **Ridge** | **Spatial** | **Hybrid** |
| CoLA (MCC) | 42.68 | 40.05 | **43.28** | **44.40** | 41.14 | 40.36 |
| SST-2 (Acc.) | 89.68 | **90.14** | 88.53 | **91.17** | **90.94** | **90.37** |
| MRPC (Acc.) | 84.80 | **84.80** | 84.10 | **86.27** | 84.56 | 84.07 |
| MRPC (F1) | 88.56 | **89.01** | **88.59** | **89.96** | **88.69** | 88.33 |
| STS-B (Pears.) | 88.06 | **88.42** | 86.14 | **88.36** | 85.84 | 87.88 |
| STS-B (Spear.) | 87.76 | **88.22** | 86.10 | **88.25** | 85.96 | **87.82** |
| QQP (Acc.) | 90.22 | **90.47** | **90.36** | **90.57** | **90.40** | **90.55** |
| QQP (F1) | 86.70 | **87.07** | **87.01** | **87.24** | **86.97** | **87.23** |
| MNLI-m (Acc.) | 82.09 | **82.66** | 81.27 | **82.33** | **82.24** | **82.31** |
| MNLI-mm (Acc.) | 82.38 | **82.97** | 82.23 | **82.98** | **82.69** | **82.83** |
| QNLI (Acc.) | 91.14 | **91.18** | **91.38** | **91.34** | **91.38** | **91.41** |
| RTE (Acc.) | 67.15 | 65.70 | **67.51** | 66.06 | **68.59** | 65.62 |
| WNLI (Acc.) | 53.52 | **56.34** | **56.34** | **56.34** | **56.34** | **57.75** |

| | | mBERT-ft-zh | | | |
|---|---|---|---|---|---|
| **CLUE Task** | **mBERT** | **MSE** | **Ridge** | **Spatial** | **Hybrid** |
| AFQMC (Acc.) | 69.74 | **71.13** | **71.78** | **70.48** | 69.51 |
| CMNLI (Acc.) | 78.66 | **78.86** | **78.79** | 78.31 | 78.87 |
| CSL (F1) | 81.10 | **81.28** | 80.83 | 80.73 | 81.43 |
| IFLYTEK (Acc.) | 56.52 | **56.87** | **56.83** | **56.71** | 56.94 |
| TNEWS (Acc.) | 54.77 | **55.00** | **55.17** | **54.91** | 55.28 |
| TNEWS (F1) | 53.69 | 53.27 | **53.89** | 52.99 | 53.61 |
| ChID (Acc.) | 10.66 | **10.66** | **10.66** | **10.66** | 10.66 |
| $C^3$ (Acc.) | 49.42 | **50.79** | **50.31** | **50.66** | 50.45 |

**b) Cross-Language Transfer Between Known Languages**

| | | mBERT-ft-zh | | | | |
|---|---|---|---|---|---|---|
| **GLUE Task** | **mBERT** | **Contrastive** | **MSE** | **Ridge** | **Spatial** | **Hybrid** |
| CoLA (MCC) | 40.96 | **41.22** | 41.83 | 41.98 | **43.31** | 42.99 |
| SST-2 (Acc.) | 89.25 | **89.25** | **90.83** | **90.14** | 89.56 | 90.94 |
| MRPC (Acc.) | 83.82 | **83.82** | **85.05** | **86.52** | **85.78** | 85.29 |
| MRPC (F1) | 87.15 | **87.88** | **88.81** | **90.02** | **89.68** | 89.29 |
| STS-B (Pears.) | 87.09 | **88.11** | 86.03 | 87.76 | 87.72 | 87.73 |
| STS-B (Spear.) | **86.97** | **87.83** | 86.18 | 87.60 | 87.58 | 87.66 |
| QQP (Acc.) | 89.91 | **90.16** | **90.42** | **90.49** | 90.14 | 90.37 |
| QQP (F1) | 86.27 | **86.89** | **87.01** | **87.11** | 86.59 | 86.93 |
| MNLI-m (Acc.) | 81.52 | **82.13** | **82.19** | **82.41** | 81.42 | 82.28 |
| MNLI-mm (Acc.) | 81.94 | **82.36** | **83.05** | **82.79** | **82.45** | 82.53 |
| QNLI (Acc.) | 90.55 | **90.92** | **91.52** | **91.18** | 90.88 | 91.16 |
| RTE (Acc.) | 66.06 | **66.43** | **67.51** | **68.95** | **70.76** | 66.79 |
| WNLI (Acc.) | 54.93 | **56.34** | **56.34** | **56.34** | **56.34** | 56.34 |

| | | mBERT-ft-en | | | |
|---|---|---|---|---|---|
| **CLUE Task** | **mBERT** | **MSE** | **Ridge** | **Spatial** | **Hybrid** |
| AFQMC (Acc.) | 69.74 | 69.07 | 69.28 | **71.78** | 71.46 |
| CMNLI (Acc.) | 78.66 | **78.68** | **78.82** | 78.48 | 78.67 |
| CSL (F1) | 81.10 | 77.92 | **81.93** | **82.07** | 80.94 |
| IFLYTEK (Acc.) | 56.52 | 56.18 | **56.52** | **57.33** | 56.18 |
| TNEWS (Acc.) | 54.77 | 54.63 | **55.27** | **54.79** | 54.92 |
| TNEWS (F1) | **53.69** | 53.01 | 53.57 | 53.43 | 53.72 |
| ChID (Acc.) | 10.66 | **10.66** | **10.66** | **10.66** | 10.66 |
| $C^3$ (Acc.) | 49.42 | 48.40 | **50.03** | **50.05** | 49.76 |

#### 2 Comparison of Multilingual Language Models

Consistency with prior work: Previous brain-informed tuning approaches (Schwartz et al., 2019; Moussa et al., 2024; Vattikonda et al., 2025) with monolingual english brain data have predominantly used encoders (e.g., BERT, Wav2Vec, Whisper, WavLM) due to their strong alignment with text and speech features. Bilingual dataset requirements: Our dataset contains bilingual brain recordings. To ensure architectural uniformity across English and Chinese experiments, we used BERT-base, BERT-Chinese, and mBERT, which share a comparable encoder backbone. However, our experiments are not limited to encoders. We also fine-tuned XLM-R (encoder) and XGLM (decoder), showing that our brain-informed fine-tuning approach generalizes across architectures Regarding model comparisons, we include fine-tuned results for mBERT, XLM-R, and XGLM, as shown in Table 2. This table shows that fine-tuning improves performance on most tasks: mBERT (7/9 tasks), XLM-R (9/9), and XGLM (6/9), with especially large gains for XLM-R (e.g., RTE: 54.15 → 62.71, MRPC: 69.61 → 77.94).

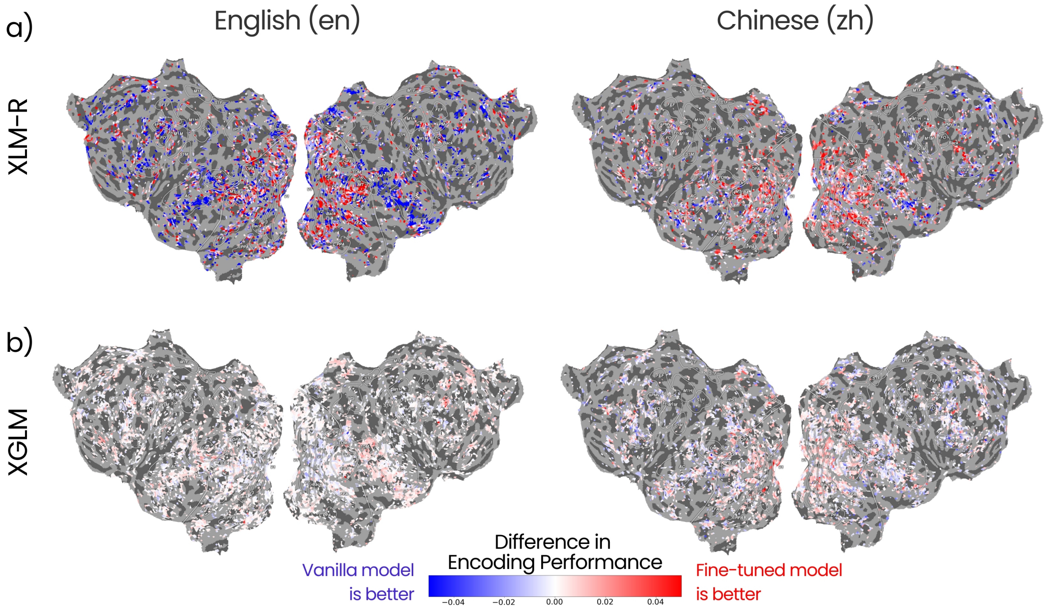


Figure 1: **Brain encoding performance before and after brain-informed fine-tuning of multilingual language models.** We evaluated the effect of bilingual brain-informed fine-tuning using voxelwise encoding models (VEMs; see Methods). VEMs were trained using representations from both the vanilla (pretrained) model and the bilingual brain-informed fine-tuned (using whole-brain) model. Flattened cortical surface for one participant is shown. Each point represents a voxel that was consistently well-predicted (r>0.1) across both models for (a) XLM-R and (b) XGLM with English (left) and Chinese (right) brain data, respectively. The VEM comparison before and after brain-informed fine-tuning shows a similar pattern to the results reported for XGLM in the main paper. However, for XLM-R, the differences look notably different. Interestingly, the fine-tuned model exhibits higher encoding performance in the early visual cortex, yet still performs well on downstream tasks. This discrepancy may arise due to spurious correlations in the stimulus or due to architectural differences in the model (cross-lingual).

#### 3 Results with LLaMA-3.2-1B

Table 3 shows downstream task performance with LLaMA-3.2-1B fine-tuned using our brain-informed approach with LoRA adaptation applied to layer 9 (similar to Antonello et al. (2024)). Models were evaluated on three downstream datasets (Wang et al., 2019; Sakaguchi et al., 2019). Brain-informed fine-tuning yields improvements over the base model across two datasets, further supporting the generalizability of our approach to different architectures and low-parameter adaptation methods (LoRA). We note that improvements are small, likely due to the parameter-efficient adaptation (LoRA on a single layer). Unfortunately, we could not perform full fine-tuning on this model due to computational constraints.

Table 2: **Comparison of downstream task performance after bilingual brain-informed fine-tuning (with whole-brain) for multilingual language models.** We compare several multilingual language models (mBERT, XLM-R, and XGLM) that undergo bilingual brain-informed fine-tuning (e.g., XLM-R-ft-en: XLM-R fine-tuned using English brain data). We evaluate downstream task performance in two settings: (a) Fine-tuning and evaluation in the same language: models are fine-tuned with brain data in one language (en) and evaluated on NLP tasks in the same language (GLUE benchmark). This tests within-language improvements due to brain-informed fine-tuning. (b) Cross-language transfer between known languages: model is fine-tuned on brain data in one language (zh) and evaluated on tasks in the participants' second (not used in fine-tuning) language (fine-tuned with zh brain data and evaluated on GLUE (en benchmark) tasks). This tests whether bilingual brain-informed fine-tuning elicits the participants' shared semantic representations. Bolded values indicate the best performance among the three fine-tuned models. Models were fine-tuned by minimizing the NT-Xent (cross-entropy) loss.

**a) Fine-tuning and Evaluation in the Same Language**

| GLUE Task | mBERT-en | mBERT-ft-en | XLM-R-en | XLM-R-ft-en | XGLM-en | XGLM-ft-en |
|---|---|---|---|---|---|---|
| CoLA (MCC) | **42.68** | 40.05 | 46.30 | **50.20** | 40.11 | 41.22 |
| SST-2 (Acc.) | 89.68 | **90.14** | 92.16 | **92.55** | 90.94 | 90.48 |
| MRPC (Acc.) | 84.80 | **84.80** | 69.61 | 77.94 | 71.32 | 71.81 |
| MRPC (F1) | 88.56 | **89.01** | 81.55 | 84.10 | 81.52 | 82.39 |
| STS-B (Pears.) | 88.06 | **88.42** | 84.96 | 85.18 | 81.43 | 82.36 |
| STS-B (Spear.) | 87.76 | **88.22** | 84.84 | 85.10 | 81.47 | 82.41 |
| QQP (Acc.) | 90.22 | 90.47 | 90.28 | **90.61** | 88.93 | 90.11 |
| QQP (F1) | 86.70 | 87.07 | 87.05 | **87.45** | 83.11 | 85.33 |
| MNLI-m (Acc.) | 82.09 | 82.66 | 84.16 | **84.46** | 76.62 | 78.85 |
| MNLI-mm (Acc.) | 82.38 | 82.97 | 84.22 | **84.40** | 78.80 | 79.01 |
| QNLI (Acc.) | 91.14 | **91.18** | 90.17 | 90.24 | 88.94 | 88.44 |
| RTE (Acc.) | **67.15** | 65.70 | 54.15 | 62.71 | 53.79 | 53.43 |
| WNLI (Acc.) | 53.52 | **56.34** | 43.66 | **56.34** | **56.34** | 56.34 |

**b) Cross-Language Transfer Between Known Languages**

| GLUE Task | mBERT-ft-zh | XLM-R-ft-zh | XGLM-ft-zh |
|---|---|---|---|
| CoLA (MCC) | 41.22 | **48.64** | 40.66 |
| SST-2 (Acc.) | 89.25 | **92.20** | 90.71 |
| MRPC (Acc.) | **83.82** | 81.86 | 73.53 |
| MRPC (F1) | **87.88** | 87.06 | 83.02 |
| STS-B (Pears.) | **88.11** | 85.90 | 82.90 |
| STS-B (Spear.) | **87.83** | 85.79 | 82.93 |
| QQP (Acc.) | 90.16 | **90.70** | 90.10 |
| QQP (F1) | 86.89 | **87.58** | 85.91 |
| MNLI-m (Acc.) | 82.13 | **84.39** | 80.55 |
| MNLI-mm (Acc.) | 82.36 | **84.85** | 80.91 |
| QNLI (Acc.) | **90.92** | 90.57 | 89.73 |
| RTE (Acc.) | **66.43** | 62.45 | 51.99 |
| WNLI (Acc.) | **56.34** | **56.34** | **56.34** |

Table 3: Downstream task performance with LLaMA-3.2-1B fine-tuned using our brain-informed approach with LoRA adaptation.

| Dataset | LLaMA-3.2-1B | LLaMA-3.2-1B-ft-en | LLaMA-3.2-1B-ft-zh |
|---|---|---|---|
| BoolQ (Acc.) | 0.5790 | **0.5848** | **0.5830** |
| COPA (Acc.) | 0.5534 | 0.5534 | 0.5534 |
| WinoGrande (Acc.) | 0.4945 | **0.4961** | **0.4961** |
| **Average** | 0.5423 | **0.5447** | **0.5441** |

### 4 Performance on Downstream NLP Tasks for Individual Participants

Table 4, 5 and 6 report downstream task performance before and after bilingual brain-informed fine-tuning (using whole-brain, language-selective, and semantically-selective voxels) across six bilingual participants.

### 5 Comparison with a Previously Proposed Fine-Tuning Pipeline

We compare our brain-informed fine-tuning approach with the pipeline used in prior work, where stimulus–TR alignment was performed before fine-tuning. Specifically, stimulus–TR pairs were constructed following the procedure described in Schwartz et al. (2019). We then fine-tuned the language model using this pre-aligned input and bilingual brain data from 1 participant. Table 8 presents downstream task performance (on the GLUE benchmark) for this preprocessing-based

Table 4: **Downstream task performance before and after bilingual brain-informed fine-tuning (with whole-brain) across subjects.** We compared vanilla (pretrained) models, English BERT (BERT-en), Chinese BERT (BERT-zh), and mBERT, with their bilingual brain-informed fine-tuned counterparts (e.g., BERT-ft-en: BERT-en fine-tuned using English brain data).

##### a) Fine-tuning and Evaluation in the Same Language

| | Participants | | | | | | | | | | | |
|---|---|---|---|---|---|---|---|---|---|---|---|---|
| **GLUE Task** | **BERT-ft-en** | | | | | | **mBERT-ft-en** | | | | | |
| | Subj1 | Subj2 | Subj3 | Subj4 | Subj5 | Subj6 | Subj1 | Subj2 | Subj3 | Subj4 | Subj5 | Subj6 |
| CoLA (MCC) | 55.75 | 54.95 | 54.16 | 55.98 | 54.69 | 56.25 | 40.05 | 41.95 | 37.64 | 37.84 | 40.40 | 37.45 |
| SST-2 (Acc.) | 93.12 | 92.74 | 92.19 | 92.66 | 92.20 | 92.55 | 90.14 | 90.71 | 90.25 | 90.60 | 91.51 | 90.14 |
| MRPC (Acc.) | 79.66 | 79.45 | 80.64 | 81.13 | 81.86 | 80.39 | 84.80 | 85.29 | 85.05 | 85.78 | 86.03 | 85.46 |
| MRPC (F1) | 86.19 | 86.08 | 87.07 | 87.40 | 87.79 | 86.97 | 89.01 | 89.01 | 89.01 | 89.38 | 89.58 | 89.16 |
| STS-B (Pears.) | 88.01 | 88.10 | 88.20 | 88.06 | 88.14 | 88.11 | 88.42 | 88.62 | 88.40 | 88.49 | 88.41 | 88.52 |
| STS-B (Spear.) | 87.63 | 87.50 | 87.71 | 87.24 | 87.62 | 87.52 | 88.22 | 88.41 | 88.23 | 88.30 | 88.30 | 88.43 |
| QQP (Acc.) | 90.77 | 90.78 | 90.84 | 90.74 | 90.79 | 90.92 | 90.47 | 90.42 | 90.42 | 90.38 | 90.47 | 90.47 |
| QQP (F1) | 87.60 | 87.64 | 88.01 | 87.57 | 87.62 | 87.85 | 87.07 | 87.08 | 87.08 | 87.05 | 87.11 | 87.12 |
| MNLI-m (Acc.) | 84.28 | 84.45 | 84.34 | 84.51 | 84.43 | 84.37 | 82.66 | 82.01 | 82.16 | 82.17 | 82.19 | 82.07 |
| MNLI-mm (Acc.) | 84.26 | 84.73 | 84.52 | 84.70 | 84.61 | 84.67 | 82.97 | 82.72 | 82.61 | 82.64 | 82.80 | 82.55 |
| QNLI (Acc.) | 91.56 | 91.59 | 91.43 | 91.38 | 91.46 | 91.34 | 91.18 | 91.31 | 91.19 | 91.23 | 91.28 | 91.02 |
| RTE (Acc.) | 67.15 | 67.34 | 67.43 | 67.26 | 67.06 | 66.82 | 65.70 | 66.06 | 64.98 | 64.92 | 67.51 | 67.51 |
| WNLI (Acc.) | 53.52 | 56.34 | 53.52 | 55.32 | 55.30 | 53.66 | 56.34 | 56.34 | 56.34 | 56.34 | 56.34 | 56.34 |

| | Participants | | | | | | | | | |
|---|---|---|---|---|---|---|---|---|---|---|
| **CLUE Task** | **BERT-ft-zh** | | | | | **mBERT-ft-zh** | | | | |
| | Subj1 | Subj2 | Subj3 | Subj4 | Subj5 | Subj1 | Subj2 | Subj3 | Subj4 | Subj5 |
| AFQMC (Acc.) | 74.01 | 74.26 | 74.37 | 75.28 | 74.56 | 70.55 | 70.72 | 69.93 | 69.96 | 72.36 |
| CMNLI (Acc.) | 80.89 | 80.82 | 80.84 | 80.92 | 80.87 | 79.02 | 79.22 | 78.87 | 78.88 | 78.96 |
| CSL (F1) | 80.73 | 81.36 | 80.44 | 80.42 | 81.13 | 80.71 | 81.62 | 81.53 | 82.11 | 82.03 |
| IFLYTEK (Acc.) | 60.25 | 60.28 | 60.56 | 60.38 | 60.87 | 56.83 | 56.91 | 57.21 | 56.84 | 56.79 |
| TNEWS (Acc.) | 56.61 | 56.49 | 56.40 | 56.46 | 56.43 | 55.12 | 54.88 | 55.30 | 55.13 | 54.96 |
| TNEWS (F1) | 55.17 | 55.36 | 55.32 | 55.34 | 55.43 | 53.74 | 53.94 | 53.76 | 53.47 | 53.45 |
| ChID (Acc.) | 10.66 | 10.66 | 10.66 | 10.66 | 10.66 | 10.66 | 10.66 | 10.66 | 10.66 | 10.66 |
| $C^3$ (F1) | 49.97 | 50.13 | 49.91 | 49.98 | 49.14 | 50.21 | 50.42 | 49.97 | 49.74 | 49.92 |

##### b) Cross-Language Transfer Between Known Languages

| | Participants | | | | | | | | | | | |
|---|---|---|---|---|---|---|---|---|---|---|---|---|
| **GLUE Task** | **BERT-ft-zh** | | | | | | **mBERT-ft-zh** | | | | | |
| | Subj1 | Subj2 | Subj3 | Subj4 | Subj5 | Subj6 | Subj1 | Subj2 | Subj3 | Subj4 | Subj5 | Subj6 |
| CoLA (MCC) | 53.82 | 52.10 | 53.40 | 55.10 | 54.70 | 53.80 | 41.22 | 39.51 | 40.30 | 42.10 | 41.81 | 41.00 |
| SST-2 (Acc.) | 92.85 | 92.61 | 92.63 | 92.95 | 92.79 | 93.27 | 89.25 | 90.14 | 90.48 | 90.94 | 90.71 | 91.17 |
| MRPC (Acc.) | 77.45 | 77.41 | 77.66 | 77.95 | 77.20 | 77.03 | 83.82 | 85.05 | 85.78 | 86.27 | 85.08 | 85.78 |
| MRPC (F1) | 84.20 | 84.23 | 84.40 | 84.60 | 84.50 | 83.27 | 87.88 | 89.20 | 89.42 | 90.00 | 89.30 | 89.49 |
| STS-B (Pears.) | 87.95 | 87.81 | 88.05 | 88.10 | 87.90 | 87.89 | 88.11 | 88.20 | 88.42 | 88.52 | 88.25 | 88.87 |
| STS-B (Spear.) | 87.65 | 87.55 | 87.60 | 87.95 | 87.60 | 87.55 | 87.83 | 88.10 | 88.30 | 88.45 | 88.14 | 88.77 |
| QQP (Acc.) | 90.76 | 90.95 | 91.03 | 90.80 | 90.78 | 90.24 | 90.16 | 90.43 | 90.48 | 90.43 | 90.44 | 90.40 |
| QQP (F1) | 87.22 | 87.73 | 87.75 | 87.82 | 87.78 | 85.02 | 86.89 | 87.04 | 87.14 | 87.08 | 87.08 | 87.04 |
| MNLI-m (Acc.) | 84.12 | 84.22 | 84.16 | 84.20 | 84.10 | 83.92 | 82.13 | 81.99 | 81.95 | 81.77 | 81.83 | 82.01 |
| MNLI-mm (Acc.) | 84.11 | 84.18 | 84.11 | 84.30 | 84.08 | 83.88 | 82.36 | 82.92 | 82.75 | 82.95 | 82.94 | 82.78 |
| QNLI (Acc.) | 91.38 | 91.47 | 91.43 | 91.37 | 91.46 | 91.17 | 90.92 | 91.54 | 91.20 | 91.17 | 91.31 | 91.09 |
| RTE (Acc.) | 67.87 | 67.80 | 67.93 | 67.97 | 67.99 | 67.66 | 66.43 | 67.15 | 68.23 | 67.87 | 67.34 | 68.95 |
| WNLI (Acc.) | 56.34 | 56.34 | 56.34 | 56.34 | 56.34 | 56.34 | 56.34 | 54.93 | 54.31 | 56.34 | 56.34 | 56.34 |

| | Participants | | | | | | | | | |
|---|---|---|---|---|---|---|---|---|---|---|
| **GLUE Task** | **BERT-ft-en** | | | | | **mBERT-ft-en** | | | | |
| | Subj1 | Subj2 | Subj3 | Subj4 | Subj5 | Subj1 | Subj2 | Subj3 | Subj4 | Subj5 |
| AFQMC (Acc.) | 69.00 | 69.00 | 69.07 | 69.00 | 69.03 | 70.62 | 70.14 | 72.38 | 70.97 | 72.13 |
| CMNLI (Acc.) | 68.62 | 68.84 | 68.67 | 68.91 | 68.87 | 78.83 | 78.81 | 78.96 | 78.82 | 78.92 |
| CSL (F1) | 71.43 | 72.91 | 71.87 | 72.87 | 72.59 | 81.30 | 81.63 | 81.37 | 81.46 | 81.79 |
| IFLYTEK (Acc.) | 47.86 | 47.90 | 46.59 | 47.52 | 47.56 | 57.14 | 56.60 | 56.57 | 57.10 | 57.18 |
| TNEWS (Acc.) | 50.92 | 51.55 | 51.41 | 51.48 | 51.51 | 54.81 | 54.91 | 54.82 | 55.23 | 55.17 |
| TNEWS (F1) | 50.60 | 50.74 | 50.69 | 50.73 | 50.64 | 53.84 | 53.77 | 53.71 | 53.76 | 53.74 |
| ChID (Acc.) | 10.66 | 10.66 | 10.66 | 10.66 | 10.66 | 10.66 | 10.66 | 10.66 | 10.66 | 10.66 |
| $C^3$ (Acc.) | 42.64 | 41.51 | 44.58 | 44.10 | 41.71 | 49.48 | 50.29 | 49.63 | 50.21 | 50.29 |

method (denoted as BERT-ft-en (Ablation)). This allows direct comparison with our proposed end-to-end brain-informed fine-tuning pipeline. We observe a substantial drop in performance for the preprocessing-based method (BERT-ft-en (Ablation)). This suggests that stimulus-TR alignment performed prior to fine-tuning procedure likely discards valuable temporal and contextual information. In contrast, our approach integrates brain alignment during fine-tuning, enabling robust downstream performance.

Table 5: **Downstream task performance before and after bilingual brain-informed fine-tuning (with language-selective voxels) across subjects.** We compared vanilla (pretrained) models, English BERT (BERT-en), Chinese BERT (BERT-zh), and mBERT, with their bilingual brain-informed fine-tuned counterparts (e.g., BERT-ft-en: BERT-en fine-tuned using English brain data).

**a) Fine-tuning and Evaluation in the Same Language**

| | **Participants** | | | | | | | | | |
|---|---|---|---|---|---|---|---|---|---|---|
| **GLUE Task** | **BERT-ft-en** | | | | | **mBERT-ft-en** | | | | |
| | Subj2 | Subj3 | Subj4 | Subj5 | Subj6 | Subj2 | Subj3 | Subj4 | Subj5 | Subj6 |
| CoLA (MCC) | 56.01 | 54.95 | 55.88 | 55.47 | 53.92 | 41.22 | 41.03 | 41.23 | 42.64 | 42.42 |
| SST-2 (Acc.) | 92.55 | 92.78 | 92.55 | 92.66 | 92.55 | 90.94 | 90.71 | 91.86 | 90.83 | 91.40 |
| MRPC (Acc.) | 80.66 | 81.86 | 81.13 | 82.60 | 80.88 | 86.76 | 85.46 | 85.82 | 85.24 | 85.54 |
| MRPC (F1) | 86.93 | 87.71 | 87.27 | 88.34 | 87.09 | 90.07 | 89.03 | 89.28 | 88.96 | 89.05 |
| STS-B (Pears.) | 87.75 | 87.67 | 87.85 | 87.38 | 87.85 | 88.70 | 87.76 | 88.35 | 88.22 | 86.36 |
| STS-B (Spear.) | 87.64 | 87.50 | 87.67 | 87.23 | 87.67 | 88.60 | 87.66 | 88.24 | 88.21 | 86.60 |
| QQP (Acc.) | 90.84 | 90.91 | 90.87 | 90.90 | 90.87 | 90.52 | 90.44 | 90.52 | 90.42 | 90.44 |
| QQP (F1) | 87.76 | 87.82 | 87.79 | 87.79 | 87.80 | 87.21 | 87.05 | 87.20 | 87.04 | 87.07 |
| MNLI-m (Acc.) | 84.56 | 84.31 | 84.39 | 83.92 | 84.27 | 82.01 | 82.03 | 82.04 | 82.42 | 82.01 |
| MNLI-mm (Acc.) | 84.62 | 84.65 | 84.85 | 84.09 | 84.78 | 82.00 | 82.53 | 82.58 | 82.79 | 82.56 |
| QNLI (Acc.) | 91.36 | 91.31 | 91.40 | 91.27 | 91.32 | 91.01 | 90.88 | 91.07 | 91.18 | 91.05 |
| RTE (Acc.) | 68.23 | 67.34 | 67.15 | 67.48 | 67.15 | 67.15 | 67.09 | 65.34 | 65.34 | 67.87 |
| WNLI (Acc.) | 52.25 | 52.21 | 53.11 | 53.66 | 53.11 | 56.34 | 57.75 | 56.34 | 56.34 | 56.34 |

| | **Participants** | | | | | | | | | | |
|---|---|---|---|---|---|---|---|---|---|---|---|
| **CLUE Task** | **BERT-ft-zh** | | | | | | **mBERT-ft-zh** | | | | |
| | Subj1 | Subj2 | Subj3 | Subj4 | Subj5 | Subj6 | Subj2 | Subj3 | Subj4 | Subj5 | Subj6 |
| AFQMC (Acc.) | 74.72 | 73.73 | 74.12 | 73.91 | 73.56 | 73.47 | 70.23 | 71.25 | 70.00 | 70.10 | 70.32 |
| CMNLI (Acc.) | 80.54 | 80.99 | 80.89 | 81.04 | 80.91 | 80.73 | 79.43 | 78.78 | 78.93 | 78.98 | 78.83 |
| CSL (F1) | 81.54 | 81.26 | 81.21 | 80.55 | 81.07 | 81.69 | 81.20 | 80.99 | 80.87 | 80.47 | 81.03 |
| IFLYTEK (Acc.) | 60.60 | 60.25 | 60.30 | 60.33 | 60.01 | 60.95 | 56.91 | 56.98 | 55.37 | 56.02 | 56.60 |
| TNEWS (Acc.) | 56.28 | 56.60 | 56.76 | 56.46 | 56.36 | 56.46 | 54.96 | 55.20 | 54.96 | 55.39 | 55.08 |
| TNEWS (F1) | 55.10 | 55.00 | 55.34 | 54.67 | 55.61 | 54.97 | 54.08 | 53.97 | 54.00 | 54.43 | 54.21 |
| ChID (Acc.) | 10.66 | 10.66 | 10.66 | 10.66 | 10.66 | 10.66 | 10.66 | 10.66 | 10.66 | 10.66 | 10.66 |
| $C^3$ (Acc.) | 50.08 | 50.16 | 49.27 | 49.82 | 49.77 | 49.53 | 50.69 | 49.95 | 50.86 | 49.71 | 49.55 |

**b) Cross-Language Transfer Between Known Languages**

| | **Participants** | | | | | | | | | |
|---|---|---|---|---|---|---|---|---|---|---|
| **GLUE Task** | **BERT-ft-zh** | | | | | **mBERT-ft-zh** | | | | |
| | Subj2 | Subj3 | Subj4 | Subj5 | Subj6 | Subj2 | Subj3 | Subj4 | Subj5 | Subj6 |
| CoLA (Acc.) | 51.84 | 51.07 | 52.38 | 52.49 | 51.95 | 43.26 | 42.72 | 42.05 | 42.31 | 42.16 |
| SST-2 (Acc.) | 92.61 | 92.33 | 92.25 | 92.37 | 92.39 | 91.06 | 90.60 | 90.89 | 90.83 | 91.06 |
| MRPC (Acc.) | 77.96 | 77.64 | 78.43 | 77.62 | 77.05 | 85.54 | 85.29 | 85.35 | 85.78 | 85.78 |
| MRPC (F1) | 85.67 | 84.57 | 84.70 | 83.72 | 83.42 | 89.29 | 89.05 | 89.21 | 89.57 | 89.61 |
| STS-B (Pears.) | 88.76 | 87.01 | 87.89 | 87.56 | 86.94 | 88.65 | 88.09 | 88.05 | 87.51 | 88.26 |
| STS-B (Spear.) | 88.39 | 88.14 | 87.66 | 87.79 | 87.44 | 88.56 | 88.36 | 87.92 | 87.48 | 88.21 |
| QQP (Acc.) | 91.24 | 90.89 | 91.09 | 91.30 | 90.54 | 90.50 | 90.48 | 90.40 | 90.37 | 90.44 |
| QQP (F1) | 87.91 | 87.69 | 87.72 | 88.21 | 87.96 | 87.14 | 87.12 | 87.04 | 86.98 | 87.09 |
| MNLI-m (Acc.) | 84.11 | 84.23 | 84.77 | 84.24 | 83.58 | 81.92 | 82.26 | 82.23 | 82.13 | 81.91 |
| MNLI-mm (Acc.) | 84.79 | 84.52 | 83.73 | 84.06 | 83.76 | 81.91 | 82.81 | 82.52 | 82.63 | 82.50 |
| QNLI (Acc.) | 91.44 | 91.31 | 91.29 | 91.41 | 91.18 | 91.07 | 91.29 | 90.98 | 91.34 | 90.92 |
| RTE (Acc.) | 66.25 | 66.73 | 67.85 | 66.21 | 67.03 | 65.34 | 66.79 | 65.62 | 68.59 | 66.79 |
| WNLI (Acc.) | 53.34 | 56.34 | 52.15 | 53.75 | 56.34 | 53.52 | 56.34 | 56.34 | 56.34 | 54.93 |

| | **Participants** | | | | | | | | | | |
|---|---|---|---|---|---|---|---|---|---|---|---|
| **CLUE Task** | **BERT-ft-en** | | | | | | **mBERT-ft-en** | | | | |
| | Subj1 | Subj2 | Subj3 | Subj4 | Subj5 | Subj6 | Subj2 | Subj3 | Subj4 | Subj5 | Subj6 |
| AFQMC (Acc.) | 69.93 | 69.00 | 69.00 | 69.00 | 69.90 | 69.00 | 70.69 | 69.76 | 70.69 | 71.27 | 71.50 |
| CMNLI (Acc.) | 68.81 | 68.96 | 67.73 | 68.60 | 68.53 | 67.40 | 78.87 | 78.92 | 78.92 | 78.95 | 78.73 |
| CSL (F1) | 71.73 | 71.88 | 71.28 | 71.89 | 71.08 | 70.89 | 81.37 | 81.02 | 81.53 | 80.87 | 81.47 |
| IFLYTEK (Acc.) | 47.74 | 46.09 | 50.09 | 47.93 | 47.17 | 49.27 | 57.14 | 56.75 | 57.21 | 56.68 | 57.33 |
| TNEWS (Acc.) | 50.79 | 50.71 | 50.76 | 50.78 | 50.78 | 50.32 | 55.03 | 55.47 | 55.51 | 55.15 | 54.99 |
| TNEWS (F1) | 50.57 | 50.56 | 50.48 | 50.56 | 50.12 | 50.56 | 54.06 | 54.01 | 54.30 | 54.13 | 54.05 |
| ChID (Acc.) | 10.66 | 10.66 | 10.66 | 10.66 | 10.66 | 10.66 | 10.66 | 10.66 | 10.66 | 10.66 | 10.66 |
| $C^3$ (Acc.) | 55.61 | 54.80 | 55.69 | 54.14 | 55.21 | 55.63 | 50.03 | 49.97 | 50.79 | 50.10 | 50.29 |

### 6 Scaling Effects on Encoding Performance in Brain-Informed Fine-Tuning

We analyzed scaling behavior on high-performing voxels using up to 6 stories ($\approx$300 TRs each). Table 9 shows that While there is a slight upward trend in encoding performance (mean $r$: 0.151 $\rightarrow$ 0.162), the effect is small relative to the standard errors, likely due to the limited dataset size. We expect clearer scaling trends to emerge with larger brain datasets (Antonello et al., 2024).

Table 6: **Downstream task performance before and after bilingual brain-informed fine-tuning (with semantic-selective voxels) across subjects.** We compared vanilla (pretrained) models, English BERT (BERT-en), Chinese BERT (BERT-zh), and mBERT, with their bilingual brain-informed fine-tuned counterparts (e.g., BERT-ft-en: BERT-en fine-tuned using English brain data).

#### a) Fine-tuning and Evaluation in the Same Language

| | Participants | | | | | | | | | |
|---|---|---|---|---|---|---|---|---|---|---|
| **GLUE Task** | **BERT-ft-en** | | | | | **mBERT-ft-en** | | | | |
| | Subj2 | Subj3 | Subj4 | Subj5 | Subj6 | Subj2 | Subj3 | Subj4 | Subj5 | Subj6 |
| CoLA (MCC) | 57.27 | 54.43 | 54.43 | 55.21 | 54.44 | 41.02 | 41.53 | 41.59 | 42.68 | 43.01 |
| SST-2 (Acc.) | 92.20 | 92.20 | 92.20 | 92.09 | 92.20 | 90.94 | 90.71 | 90.83 | 90.83 | 90.37 |
| MRPC (Acc.) | 80.06 | 80.15 | 80.15 | 80.01 | 81.37 | 84.87 | 85.54 | 85.80 | 85.78 | 84.32 |
| MRPC (F1) | 87.04 | 87.30 | 87.27 | 87.01 | 87.58 | 88.49 | 89.45 | 89.60 | 89.61 | 88.53 |
| STS-B (Pears.) | 87.69 | 87.50 | 87.49 | 86.85 | 87.03 | 88.44 | 88.57 | 87.19 | 88.60 | 88.33 |
| STS-B (Spear.) | 87.53 | 87.34 | 87.34 | 86.84 | 87.02 | 88.39 | 88.37 | 87.06 | 88.49 | 88.30 |
| QQP (Acc.) | 90.52 | 90.81 | 90.82 | 90.75 | 90.88 | 90.37 | 90.31 | 90.49 | 90.46 | 90.58 |
| QQP (F1) | 87.21 | 87.71 | 87.73 | 87.57 | 87.79 | 87.08 | 87.01 | 87.14 | 87.08 | 87.27 |
| MNLI-m (Acc.) | 84.15 | 84.15 | 84.32 | 84.55 | 84.38 | 81.92 | 81.98 | 82.08 | 81.96 | 82.26 |
| MNLI-mm (Acc.) | 84.23 | 84.27 | 84.72 | 84.37 | 84.30 | 82.73 | 82.54 | 82.82 | 82.88 | 82.73 |
| QNLI (Acc.) | 91.54 | 91.45 | 91.61 | 91.56 | 91.40 | 91.25 | 91.20 | 91.24 | 91.05 | 91.17 |
| RTE (Acc.) | 67.15 | 67.87 | 67.87 | 63.90 | 67.06 | 68.95 | 67.79 | 68.23 | 67.15 | 68.59 |
| WNLI (Acc.) | 53.38 | 53.52 | 53.52 | 53.52 | 53.66 | 54.93 | 56.34 | 56.34 | 56.34 | 56.34 |

| | Participants | | | | | | | | | |
|---|---|---|---|---|---|---|---|---|---|---|
| **CLUE Task** | **BERT-ft-zh** | | | | | **mBERT-ft-zh** | | | | |
| | Subj2 | Subj3 | Subj4 | Subj5 | Subj6 | Subj2 | Subj3 | Subj4 | Subj5 | Subj6 |
| AFQMC (Acc.) | 74.03 | 73.66 | 73.80 | 73.15 | 74.14 | 70.09 | 70.25 | 70.42 | 70.11 | 70.23 |
| CMNLI (Acc.) | 80.60 | 81.05 | 80.95 | 81.10 | 80.97 | 79.70 | 79.05 | 79.20 | 79.25 | 79.10 |
| CSL (F1) | 81.15 | 81.37 | 81.13 | 81.39 | 82.03 | 81.58 | 80.58 | 82.20 | 81.69 | 80.84 |
| IFLYTEK (Acc.) | 60.37 | 61.00 | 59.74 | 60.69 | 60.05 | 56.75 | 56.33 | 56.60 | 56.68 | 56.68 |
| TNEWS (Acc.) | 56.32 | 56.41 | 56.50 | 56.59 | 56.68 | 55.17 | 55.02 | 55.04 | 54.78 | 55.23 |
| TNEWS (F1) | 56.12 | 56.29 | 55.61 | 55.78 | 55.95 | 54.36 | 53.66 | 54.05 | 54.21 | 54.16 |
| ChID (Acc.) | 10.66 | 10.66 | 10.66 | 10.66 | 10.66 | 10.66 | 10.66 | 10.66 | 10.66 | 10.66 |
| $C^3$ (Acc.) | 50.19 | 49.08 | 50.00 | 49.85 | 49.82 | 49.69 | 52.10 | 50.11 | 51.05 | 49.71 |

#### b) Cross-Language Transfer Between Known Languages

| | Participants | | | | | | | | | |
|---|---|---|---|---|---|---|---|---|---|---|
| **GLUE Task** | **BERT-ft-zh** | | | | | **mBERT-ft-zh** | | | | |
| | Subj2 | Subj3 | Subj4 | Subj5 | Subj6 | Subj2 | Subj3 | Subj4 | Subj5 | Subj6 |
| CoLA (MCC) | 52.62 | 51.85 | 53.16 | 53.27 | 52.73 | 41.73 | 40.14 | 41.09 | 41.17 | 41.78 |
| SST-2 (Acc.) | 92.81 | 92.53 | 92.45 | 92.57 | 92.59 | 90.94 | 90.71 | 90.14 | 90.25 | 90.51 |
| MRPC (Acc.) | 78.92 | 78.60 | 78.39 | 79.58 | 79.01 | 86.03 | 85.54 | 85.60 | 86.03 | 85.80 |
| MRPC (F1) | 87.20 | 86.10 | 85.23 | 85.10 | 84.95 | 89.80 | 89.45 | 88.68 | 89.88 | 89.45 |
| STS-B (P) | 88.30 | 88.55 | 88.43 | 88.10 | 88.48 | 88.51 | 88.32 | 88.17 | 87.35 | 88.09 |
| STS-B (SP) | 88.15 | 88.30 | 88.22 | 87.95 | 88.20 | 88.45 | 88.22 | 88.08 | 87.18 | 88.00 |
| QQP (Acc.) | 91.40 | 90.85 | 91.15 | 91.26 | 90.50 | 90.35 | 90.49 | 90.40 | 90.55 | 90.45 |
| QQP (F1) | 87.85 | 87.63 | 88.06 | 88.15 | 87.90 | 86.96 | 87.13 | 87.01 | 87.20 | 87.08 |
| MNLI-m (Acc.) | 84.51 | 84.63 | 85.17 | 84.64 | 83.98 | 81.96 | 82.04 | 82.21 | 82.07 | 82.07 |
| MNLI-mm (Acc.) | 84.17 | 84.90 | 84.11 | 84.44 | 83.64 | 82.73 | 82.72 | 82.92 | 82.58 | 82.74 |
| QNLI (Acc.) | 91.35 | 91.42 | 91.30 | 91.38 | 91.29 | 91.10 | 91.07 | 90.87 | 91.12 | 91.04 |
| RTE (Acc.) | 67.72 | 66.20 | 66.32 | 67.20 | 66.50 | 70.40 | 68.43 | 68.18 | 68.23 | 68.31 |
| WNLI (Acc.) | 56.34 | 56.34 | 56.34 | 56.34 | 56.34 | 56.34 | 56.34 | 56.34 | 56.34 | 56.34 |

| | Participants | | | | | | | | | |
|---|---|---|---|---|---|---|---|---|---|---|
| **CLUE Task** | **BERT-ft-en** | | | | | **mBERT-ft-en** | | | | |
| | Subj2 | Subj3 | Subj4 | Subj5 | Subj6 | Subj2 | Subj3 | Subj4 | Subj5 | Subj6 |
| AFQMC (Acc.) | 69.19 | 69.00 | 69.44 | 69.91 | 69.00 | 70.48 | 72.06 | 71.01 | 71.06 | 71.32 |
| CMNLI (Acc.) | 68.80 | 68.55 | 68.27 | 68.73 | 68.14 | 79.27 | 79.01 | 79.09 | 79.17 | 78.91 |
| CSL (F1) | 71.93 | 69.63 | 73.92 | 71.17 | 71.50 | 80.77 | 81.17 | 82.19 | 81.19 | 81.13 |
| IFLYTEK (Acc.) | 47.71 | 47.71 | 47.13 | 47.74 | 47.36 | 57.71 | 57.29 | 57.37 | 56.94 | 57.64 |
| TNEWS (Acc.) | 51.58 | 51.20 | 50.79 | 51.86 | 51.98 | 55.24 | 55.29 | 55.15 | 55.28 | 55.15 |
| TNEWS (F1) | 51.36 | 51.15 | 49.15 | 51.08 | 51.14 | 54.11 | 54.22 | 53.98 | 54.19 | 53.58 |
| ChID (Acc.) | 10.66 | 10.66 | 10.66 | 10.66 | 10.66 | 10.66 | 10.66 | 10.66 | 10.66 | 10.66 |
| $C^3$ (Acc.) | 50.08 | 35.08 | 45.18 | 38.94 | 35.98 | 50.79 | 50.81 | 50.13 | 50.45 | 50.03 |

Table 7: **Downstream task performance before and after bilingual or monolingual brain-informed fine-tuning.** We perform brain-informed fine-tuning of mBERT with English whole-brain brain data (mBERT-ft-en) from either a bilingual (participant 1) or a monolingual participant. We evaluate downstream task performance in two settings: (a) Fine-tuning and evaluation in the same language: the model is evaluated in English with GLUE tasks. (b) Cross-language transfer between known languages: the model is evaluated in Chinese with CLUE tasks.

**a) Fine-tuning and Evaluation in the Same Language**

| | | mBERT-ft-en | | | |
|---|---|---|---|---|---|
| **Task** | **mBERT** | **Bilingual** | **Mono1** | **Mono2** | **Mono3** |
| CoLA (MCC) | **42.68** | 40.05 | 35.74 | 46.36 | 37.96 |
| SST-2 (Acc.) | 89.68 | 90.14 | **90.48** | 90.83 | 91.97 |
| MRPC (Acc.) | 84.80 | **84.80** | 78.85 | 80.91 | 80.84 |
| MRPC (F1) | 88.56 | **89.01** | 85.78 | 86.54 | 86.03 |
| STS-B (Pearson) | 88.06 | **88.42** | **88.42** | 88.09 | 87.35 |
| STS-B (Spearman) | 87.76 | 88.22 | **88.34** | 87.85 | 87.30 |
| QQP (Acc.) | 90.22 | 90.47 | **90.54** | 90.33 | 90.42 |
| QQP (F1) | 86.70 | 87.07 | **87.19** | 86.84 | 87.05 |
| MNLI-m (Acc.) | 82.09 | **82.66** | 82.41 | 81.90 | 82.17 |
| MNLI-mm (Acc.) | 82.38 | **82.97** | 82.78 | 82.56 | 82.71 |
| QNLI (Acc.) | 91.14 | **91.18** | 91.12 | 91.14 | 91.21 |
| RTE (Acc.) | 67.15 | **65.70** | 65.34 | 65.34 | 65.34 |
| WNLI (Acc.) | 53.52 | **56.34** | **56.34** | 56.34 | 56.34 |

**b) Cross-Language Transfer Between Known Languages**

| | | mBERT-ft-en | | | |
|---|---|---|---|---|---|
| **Task** | **mBERT** | **Bilingual** | **Mono1** | **Mono2** | **Mono3** |
| AFQMC (Acc.) | 69.74 | 70.62 | **70.64** | 69.78 | 70.57 |
| CMNLI (Acc.) | 78.66 | **78.83** | 78.48 | 78.72 | 78.60 |
| CSL (F1) | 81.10 | **81.30** | 81.18 | 81.13 | 80.97 |
| IFLYTEK (Acc.) | 46.79 | 56.52 | **57.06** | 57.10 | 56.44 |
| TNEWS (Acc.) | 54.77 | **54.81** | 54.80 | 54.26 | 54.77 |
| TNEWS (F1) | 53.69 | **53.84** | 53.50 | 53.33 | 53.44 |
| ChID (Acc.) | 10.66 | **10.66** | **10.66** | **10.66** | **10.66** |
| $C^3$ (F1) | 49.42 | **49.48** | 49.46 | 49.34 | 49.04 |

Table 8: GLUE results: Baseline BERT-en, fine-tuned BERT-ft-en (mean ± std across subjects), and BERT-ft-en ablation scores.

| GLUE Task | BERT-en | BERT-ft-en (Mean ± Std) | BERT-ft-en (Ablation) |
|---|---|---|---|
| CoLA (MCC) | 53.38 | **55.11** ± **0.75** | 0.00 |
| SST-2 (Acc.) | 92.08 | **92.58** ± **0.39** | 50.92 |
| MRPC (Acc.) | 79.41 | **80.55** ± **1.01** | 68.38 |
| MRPC (F1) | 86.27 | **86.91** ± **0.75** | 81.22 |
| STS-B (Pears.) | 88.06 | **88.10** ± **0.07** | 0.008 |
| STS-B (Spear.) | **87.65** | 87.55 ± 0.18 | 0.028 |
| QQP (Acc.) | 90.84 | **90.79** ± **0.03** | 75.28 |
| QQP (F1) | 87.70 | **87.71** ± **0.19** | 62.81 |
| MNLI-m (Acc.) | 84.38 | **84.40** ± **0.09** | 41.74 |
| MNLI-mm (Acc.) | **84.64** | 84.55 ± 0.20 | 41.48 |
| QNLI (Acc.) | 91.45 | **91.49** ± **0.09** | 58.32 |
| RTE (Acc.) | 67.15 | **67.32** ± **0.17** | 52.71 |
| WNLI (Acc.) | 49.30 | **55.59** ± **1.31** | 43.66 |

Table 9: Scaling effects on encoding performance in brain-informed fine-tuning.

| **No. of Stories used for fine-tuning** | **1** | **2** | **3** | **4** | **5** | **6** |
|---|---|---|---|---|---|---|
| **Mean (r)** | 0.1515 | 0.1508 | 0.1500 | 0.1571 | 0.1624 | 0.1625 |
| **SE** | 0.0052 | 0.0054 | 0.0053 | 0.0102 | 0.0149 | 0.0150 |

Omer Moussa, Dietrich Klakow, and Mariya Toneva. Improving semantic understanding in speech language models via brain-tuning. *arXiv preprint arXiv:2410.09230*, 2024.

Keisuke Sakaguchi, Ronan Le Bras, Chandra Bhagavatula, and Yejin Choi. An adversarial winograd schema challenge at scale. *arXiv preprint arXiv:1907.10641*, 6, 2019.

Dan Schwartz, Mariya Toneva, and Leila Wehbe. Inducing brain-relevant bias in natural language processing models. *Advances in Neural Information Processing Systems*, 32, 2019.

Nishitha Vattikonda, Aditya R Vaidya, Richard J Antonello, and Alexander G Huth. Brain-wavlm: Fine-tuning speech representations with brain responses to language. *arXiv preprint arXiv:2502.08866*, 2025.

Alex Wang, Yada Pruksachatkun, Nikita Nangia, Amanpreet Singh, Julian Michael, Felix Hill, Omer Levy, and Samuel Bowman. Superglue: A stickier benchmark for general-purpose language understanding systems. *Advances in neural information processing systems*, 32, 2019.
